# Gut bacteria convert cortisol into a distinct host signalling molecule

**DOI:** 10.1101/2020.06.12.149468

**Authors:** Maayan Freiberg, Taojun Wang, Raissa Santos de Lima Rosa, Annika Osswald, Marcelo C.R. Melo, David G. Stouffer, Pius Sarfo Buobu, Caitlin Welsh, Lufan Yang, Ahmed M. Abdel-Hamid, Heidi L. Doden, Saravanan Devendran, Rebecca M. Pollet, Sarah Goldberg, Lucille Ray, Alyshia Hamm, Nicollette Kessee, Hanchu Dai, Sean M. Mythen, Shiva Bhowmik, Scott A. Lesley, Phillip B. Hylemon, Zaida Luthey-Schulten, Fuming Pan, Ece Mutlu, Steven L. Daniel, Yong-Su Jin, Sage J. Kim, Paul Grippo, Richard J. Auchus, Isaac Cann, Andrew J. Steelman, Michael Robben, Chris Greening, Nicole M. Koropatkin, Soeren Ocvirk, H. Rex Gaskins, Patricia G. Wolf, Lisa Tussing-Humphreys, Jae Won Lee, Rafael C. Bernardi, Jason M. Ridlon

**Author notes:** These authors jointly supervised this work (Maayan Freiberg, Taojun Wang, Patricia G. Wolf, Lisa Tussing-Humphreys, Jae Won Lee, Rafael C. Bernardi, Jason M. Ridlon).

## Abstract

Microorganisms extensively modify host steroids, but whether these reactions merely eliminate hormones or create signals with new biological identities is largely unknown. Gut bacteria have been known for more than four decades to reduce cortisol to 20***α***-dihydrocortisol, yet the physiological consequence of this transformation remained unresolved. Here we show that microbial cortisol reduction creates a distinct host signalling molecule. A 2.0-Å structure of the bacterial enzyme DesC, together with molecular dynamics, biochemical perturbation and hybrid quantum mechanics/molecular mechanics simulations, defines substrate recognition and an ordered hydride-transfer and proton-relay mechanism. In gnotobiotic mice, isogenic bacteria expressing active DesC—but not a catalytically inactive S47A variant—produced 20***α***-dihydrocortisol in the intestine and circulation and reprogrammed colonic transcription. In primary intestinal epithelial cells, 20***α***-dihydrocortisol, but not cortisol, activated ERK-dependent inflammatory and growth-associated programmes. These responses required nuclear receptor subfamily 4 group A member 3 (NR4A3), whose purified ligand-binding domain bound 20***α***-dihydrocortisol but showed no detectable binding to cortisol. A Chicago colonoscopy cohort linked chronic cortisol exposure, faecal *desC*, the microbial metabolite and colorectal phenotypes. Thus, bacterial metabolism can change receptor selectivity and biological activity rather than simply terminate host hormone action, expanding the endocrine chemistry of the host.

## Introduction

Steroid hormones are defined not only by their carbon skeletons but also by the placement and stereochemistry of individual functional groups. Peripheral metabolism can therefore terminate hormone action, change receptor selectivity or generate molecules with new biological activities; several host steroid-metabolizing enzymes are drug targets for precisely this reason [1]. Host-associated microorganisms add a largely unexplored layer to this chemistry. They transform cholesterol, bile acids, progestins, glucocorticoids, androgens and oestrogens, thereby extending the steroid repertoire encountered by host tissues [2–5]. Microbial bile-acid metabolism has established that bacterial products can engage host receptors and reshape metabolism and immunity [6–8]. Yet most microbial transformations of host hormones are still interpreted as degradation or clearance rather than as potential signal-generating reactions.

The gut bacterium *Clostridium scindens* carries the cortisol-inducible *desABCD* locus. DesAB cleaves the cortisol side chain, whereas the NADH-dependent 20*α*-hydroxysteroid dehydrogenase DesC reduces its C20 carbonyl to form 20*α*-dihydrocortisol [9–11]. Functional metagenomics and phylogenetic analyses have identified *desC* as the only currently characterized microbial gene with this activity [9, 12]; *C. scindens* strains nevertheless vary in steroid-metabolism gene content [13]. The reaction itself has been known for more than four decades, but whether it changes host physiology has remained unknown.

Here we connect the atomic mechanism of DesC to the biological identity of its product. Crystallography, classical molecular dynamics (MD), mutagenesis and hybrid quantum mechanics/molecular mechanics (QM/MM) simulations define how DesC recognizes cortisol and performs stereospecific reduction. The resulting catalytic-null S47A variant enables an isogenic gnotobiotic experiment showing that bacterial DesC produces circulating 20*α*-dihydrocortisol and changes intestinal transcription. We further identify 20*α*-dihydrocortisol as a ligand of NR4A3 that elicits a cellular programme distinct from cortisol. Finally, a human cohort establishes that the enzyme, metabolite and relevant host exposures occur together in a population with colorectal phenotyping. The core finding is therefore not microbial hormone disposal, but microbial creation of a host signal.

## Results

### Atomistic mechanism of microbial cortisol reduction

We first defined the chemical selectivity and structure of DesC (Fig. 1a–c). Recombinant DesC reduced cortisol and several closely related glucocorticoids, but did not act detectably on corticosterone, dexamethasone, progesterone, allopregnanolone or aldosterone. This profile differed from that of human 20*α*-hydroxysteroid dehydrogenase (AKR1C1), whose preferred substrate was progesterone and whose activity across the corticosteroid panel was distinct (Extended Data Fig. 1). The bacterial enzyme therefore recognizes a restricted combination of steroid substituents rather than functioning as a general steroid reductase.

**Fig. 1.**
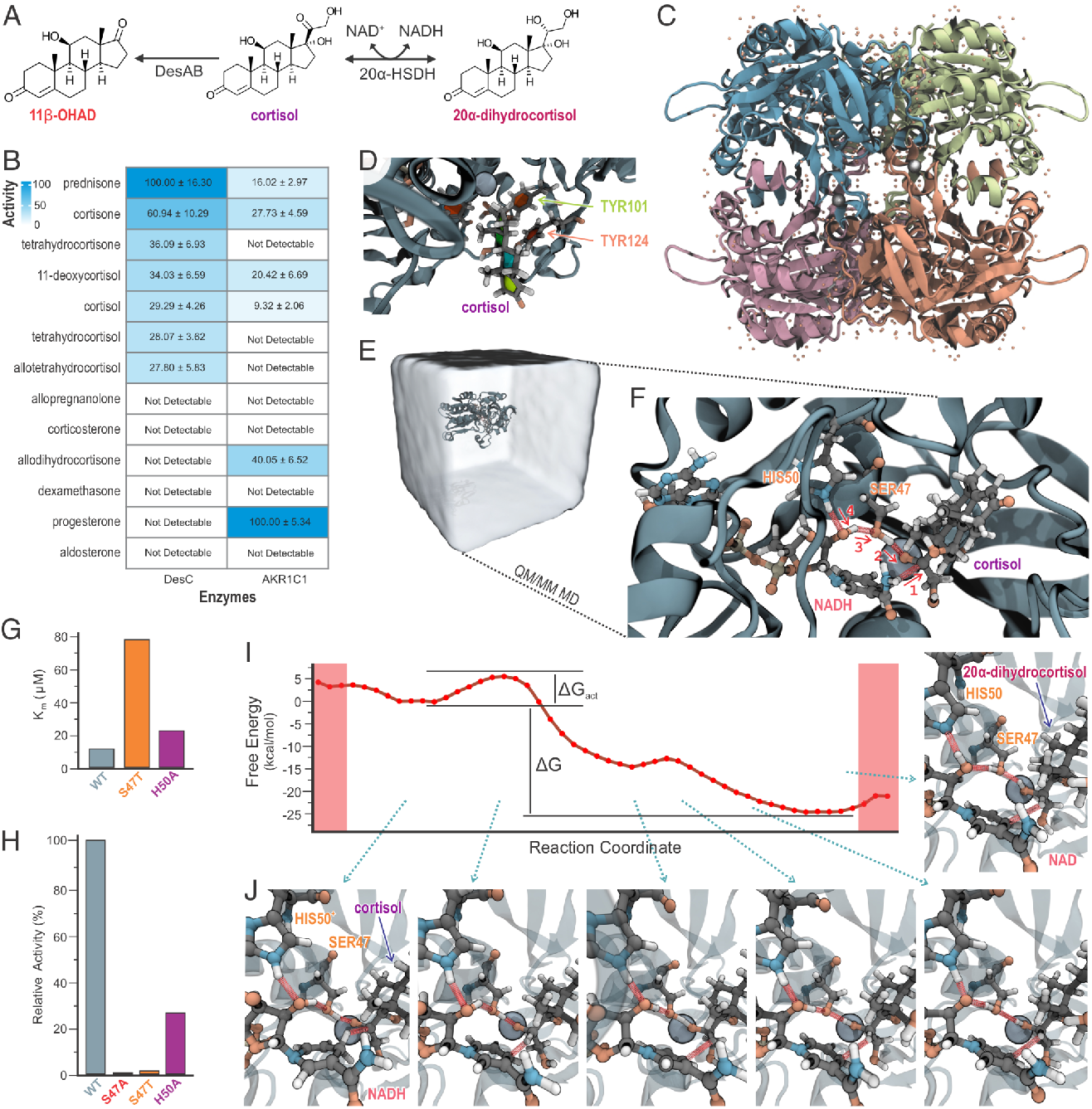
DesC recognizes cortisol and reduces its C20 carbonyl through an ordered chemical pathway. **a**, The cortisol-inducible *des* pathway. DesAB cleaves the cortisol side chain to form 11*β*-hydroxyandrostenedione (11*β*-OHAD; red), whereas the NADH-dependent 20*α*-hydroxysteroid dehydrogenase DesC reduces cortisol (purple) to 20*α*-dihydrocortisol (pink). **b**, Relative activity of recombinant DesC and human 20*α*-HSDH (AKR1C1) with corticosteroids and neurosteroids (200 *µ*M). Values were normalized separately to prednisone for DesC and progesterone for AKR1C1 and are mean*±*s.d. from *n* = 3 biological replicates per substrate; not detectable indicates no activity above the assay threshold. **c**, Tetrameric 2.0-Å apo-DesC structure (PDB 4OH1). Monomers are coloured separately, structural waters are orange spheres and zinc ions are grey spheres. **d**, MD-derived cortisol pose showing Tyr101 (green) and Tyr124 (salmon) in the hydrophobic recognition pocket. **e**, Explicit-solvent DesC system used for classical and QM/MM simulations. **f**, QM/MM active site and the four numbered bond rearrangements. Hydride transfers from NADH to the carbon atom of the cortisol C20 carbonyl; Ser47 protonates the carbonyl oxygen; transfers from the NAD^+^ ribose to Ser47 and from His50 to the ribose then restore the relay. **g,h**, Cortisol Michaelis constant (**g**) and relative activity (**h**) of wild-type (WT) DesC and the S47A, S47T and H50A variants under saturating cofactor and substrate conditions. S47A had no measurable activity and is absent from **g**; bars are mean*±*s.d. from *n* = 3 biological replicates. **i**, PM7 QM/MM free-energy profile obtained with multiple-walker extended adaptive biasing force along the optimized string. Δ*G*_act_ and Δ*G* describe the selected chemical coordinate. **j**, Active-site configurations at the five indicated positions along the pathway. The calculation used 750 string replicas and 600 free-energy walkers.

The apo structure, resolved to 2.0 Å, revealed a zinc-dependent medium-chain dehydrogenase tetramer with a Rossmann-fold nucleotide-binding domain and a catalytic domain (Fig. 1c; PDB 4OH1) [14, 15]. Electron-density and coordination analyses define its catalytic and structural zinc sites (Extended Data Fig. 2), while comparison with cofactor-bound homologues provided the framework for NADH placement (Extended Data Fig. 3). Because the crystal lacked substrates, we positioned cortisol with its C20 carbonyl adjacent to the nicotinamide hydride donor and catalytic zinc before 100 ns of explicit-solvent MD [16–18]. The model predicted that Tyr101 and Tyr124 stabilize cortisol in a hydrophobic pocket (Fig. 1d). This prediction was experimentally discriminating: Y101R and Y124R retained their overall secondary structure but lost 96.0*±*0.7% and 81.8*±*0.3% of catalytic activity, respectively, with severely impaired cortisol recognition (Extended Data Fig. 4). Thus, the simulations identified substrate-recognition residues not apparent from the apo structure alone.

We next resolved the chemical pathway using electrostatically embedded QM/MM MD with a 573-atom quantum region (Fig. 1e,f) [19, 20]. The favoured pathway contains no intervening water: hydride transfers from NADH to the C20 carbonyl carbon while Ser47 protonates the carbonyl oxygen. The catalytic state is then restored through a relay from the NAD^+^ ribose to Ser47 and from protonated His50 to the ribose. S47A abolished activity while preserving cortisol binding, whereas S47T lost both nearly all activity and detectable cortisol binding; H50A retained partial activity (Fig. 1g,h; Supplementary Tables 2c,d). These perturbations distinguish catalytic chemistry from gross protein destabilization and support the roles assigned computationally.

We optimized the reaction coordinate with 750 string-method replicas and sampled its free-energy profile with 600 multiple-walker extended adaptive-biasing-force calculations [21–23]. The optimized order was hydride transfer, protonation of the nascent alkoxide and restoration of the proton relay (Fig. 1i,j; Extended Data Fig. 5). Within the PM7 QM/MM model, hydride transfer produced the resolved chemical barrier and the subsequent proton transfers were downhill. Together, the experiments and calculations define an atomically explicit mechanism and, critically, provide the catalytically inactive S47A control required to isolate DesC chemistry in vivo.

### DesC creates a circulating microbial cortisol product

Because *C. scindens* remains difficult to manipulate genetically [24, 25], we integrated either wild-type *desC* or *desC* S47A into the same intergenic locus of *Escherichia coli*, under the same constitutive promoter (Fig. 2a). Reporter validation, chromosomal insertion and plasmid-curing controls established stable expression from this locus (Extended Data Fig. 6). The resulting Eco_desCWT and Eco_desCM strains had comparable growth, but only Eco_desCWT converted cortisol to 20*α*-dihydrocortisol, reaching 57.9*±*1.2% conversion in culture (Fig. 2b). The design therefore holds bacterial background and expression architecture constant while changing catalytic turnover.

**Fig. 2.**
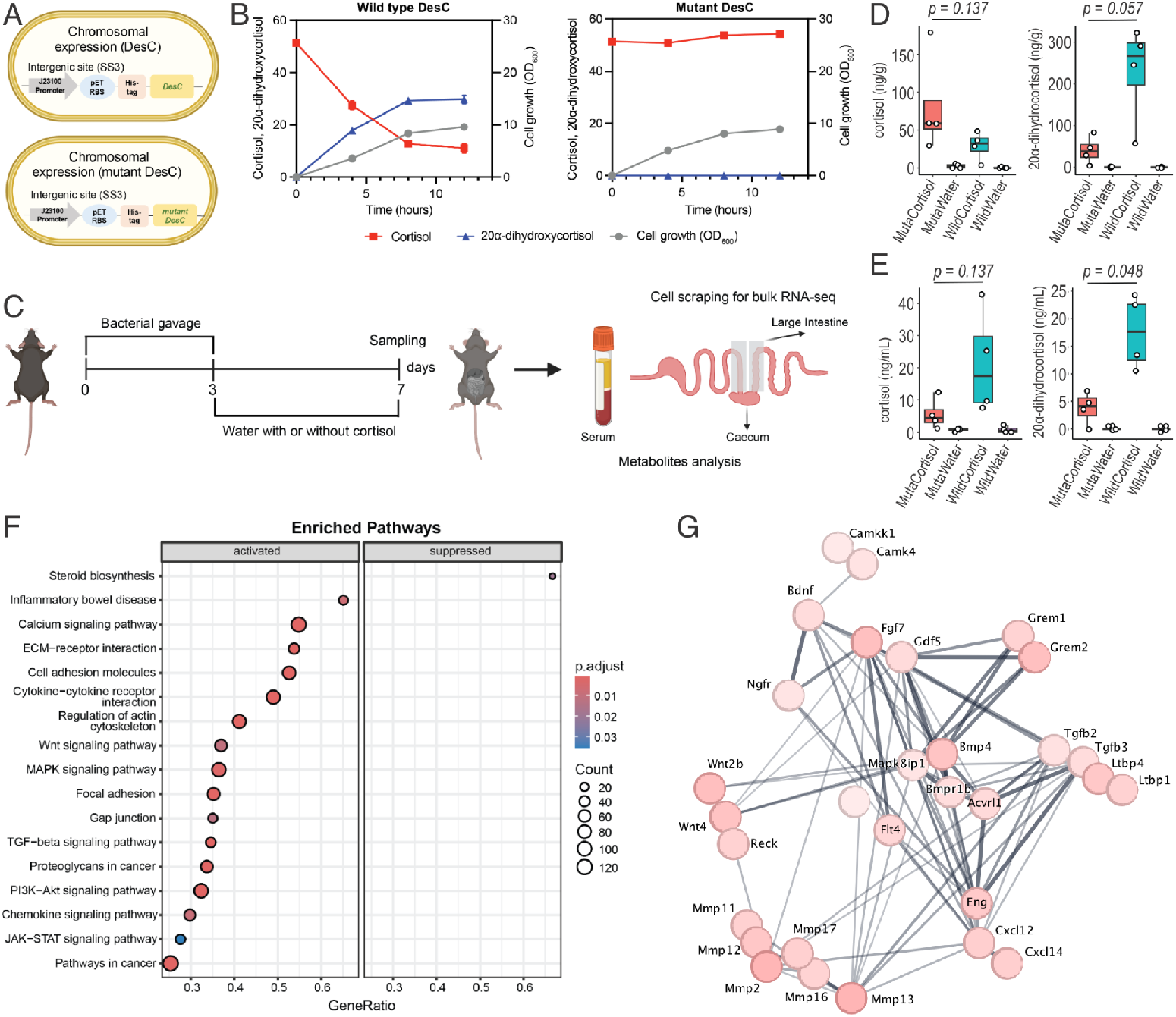
Bacterial DesC produces circulating 20*α*-dihydrocortisol and reprogrammes intestinal transcription. **a**, Isogenic chromosomal cassettes expressing His-tagged wild-type DesC or catalytically inactive DesC S47A from the J23100 promoter and pET ribosome-binding site at the SS3 intergenic locus of *E. coli*. **b**, Cortisol depletion (red squares), 20*α*-dihydrocortisol production (blue triangles) and bacterial growth (grey circles; OD_600_) over 12 h. Wild-type DesC converted cortisol, whereas mutant DesC did not; points and error bars show mean*±*s.d. from independent cultures. **c**, Gnotobiotic design. Germ-free C57BL/6J mice (n=4/group) were gavaged with engineered bacteria (10^8^ CFU/gavage) on days 1 and 3, given sterile water with or without cortisol (10 mg kg*^−^*^1^ day*^−^*^1^) from days 4–7, and sampled on day 7 for serum and caecal steroid analysis and large-intestinal RNA sequencing. **d,e**, Cortisol and 20*α*-dihydrocortisol in caecal contents (**d**) and serum (**e**). MutaCorti-sol and MutaWater denote mice carrying S47A DesC and receiving cortisol or water; WildCortisol and WildWater denote mice carrying wild-type DesC and receiving cortisol or water. Salmon and turquoise boxes denote the mutant-and wild-type-DesC cortisol groups, respectively; water groups are unfilled. Each point is one mouse (*n* = 4 analysed mice per group). Boxes show medians and interquartile ranges; whiskers extend to 1.5 times the interquartile range. Indicated *P* values are from two-sided Wilcoxon rank-sum tests with Benjamini–Hochberg correction. **f**, KEGG enrichment among differentially expressed genes in large-intestinal epithelial scrapings from WildCortisol versus MutaCortisol mice (*n* = 4 biological replicates per group). Point size indicates gene count, horizontal position indicates gene ratio and colour indicates the adjusted *P* value. **g**, Functional interaction network for representative induced genes from the same contrast; nodes are genes and edges are curated or predicted functional associations.

Germ-free C57BL/6J mice were mono-associated with either strain and then given water with or without cortisol (four analysed mice per group; Fig. 2c). Mice do not synthesize cortisol endogenously [26], allowing the administered substrate to be tracked against the murine glucocorticoid background. Mono-association was confirmed by 16S rRNA gene sequencing (Extended Data Fig. 7). Cortisol exposure was similar between colonization groups, whereas 20*α*-dihydrocortisol accumulated in the caecum and was significantly higher in serum from mice carrying active DesC than in mice carrying S47A DesC (Fig. 2d,e). A single bacterial carbonyl-reduction reaction in the gut can therefore generate a circulating steroid.

Large intestinal epithelial scrapings from cortisol-treated mice carrying Eco_desCWT or Eco_desCM were then compared by RNA sequencing. Active DesC induced 857 genes and enriched MAPK, WNT, calcium, focal-adhesion, inflammatory and cancer-associated programmes (Fig. 2f,g). The catalytic state of one bacterial enzyme is therefore sufficient to determine whether a cortisol-derived signal reaches the circulation and changes host intestinal transcription. These data do not demonstrate tumour formation; rather, they show that the microbial product reaches host tissues and changes intestinal programmes that control signalling, inflammation and growth.

### The microbial product engages NR4A3

To separate the effect of the metabolite from other consequences of bacterial colonization, we exposed primary mouse intestinal epithelial cells to 500 nM 20*α*-dihydrocortisol, cortisol or vehicle. The microbial product upregulated 91 genes and downregulated 97 genes, with enrichment of MAPK, TNF, NF-*κ*B, JAK–STAT, focal-adhesion and cancer-associated programmes and suppression of cell-cycle, DNA-replication and repair programmes (Fig. 3a; Extended Data Fig. 8). Cortisol did not reproduce this transcriptional pattern. At the protein level, 20*α*-dihydrocortisol, but not cortisol, increased the pERK/ERK ratio and nuclear pERK after six hours (Fig. 3b,c). Thus, bacterial reduction creates biological activity distinct from the parent glucocorticoid. The absence of an acute 20-min response (Extended Data Fig. 7) was more consistent with an intracellular transcriptional mechanism than with immediate activation of a cell-surface receptor.

**Fig. 3.**
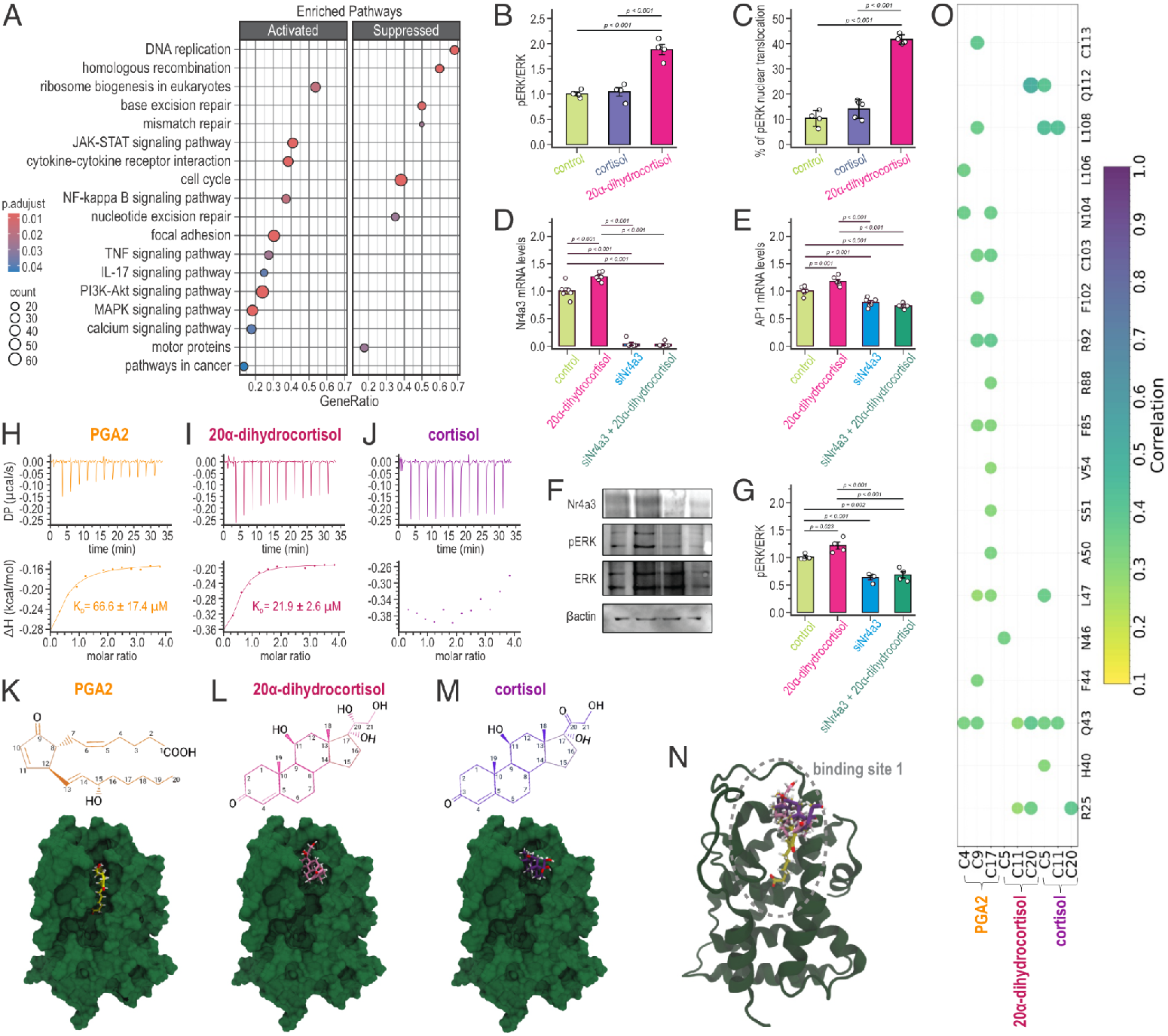
20α-dihydrocortisol engages NR4A3 and activates ERK signalling in intestinal epithelial cells. **a**, KEGG enrichment among genes altered after 6 h exposure of primary mouse intestinal epithelial cells to 500 nM 20*α*-dihydrocortisol versus vehicle (*n* = 4 biological replicates). Point size indicates gene count, horizontal position indicates gene ratio and colour indicates the Benjamini–Hochberg-adjusted *P* value. **b,c**, pERK/ERK ratio (**b**) and percentage of cells with nuclear pERK (**c**) after vehicle (light green), cortisol (violet) or 20*α*-dihydrocortisol (pink) treatment (*n* = 4 biological replicates). **d,e**, *Nr4a3* (**d**) and AP-1-complex (**e**) mRNA after control siRNA with vehicle (light green), control siRNA with 20*α*-dihydrocortisol (pink), *Nr4a3* siRNA (blue), or *Nr4a3* siRNA with metabolite (green); each point is one biological replicate (*n* = 4–6 per group). **f**, Representative immunoblots for NR4A3, pERK, ERK and *β*-actin under the same four conditions. **g**, pERK/ERK quantification (*n* = 4 biological replicates). Bars in **b–e,g** show mean*±*s.d.; indicated *P* values are from two-sided unpaired *t*-tests with Benjamini–Hochberg correction. **h–j**, Representative ITC thermograms and fitted isotherms for purified mouse NR4A3 ligand-binding domain with PGA2 (orange; **h**), 20*α*-dihydrocortisol (pink; **i**) or cortisol (purple; **j**). *K*_D_ = 66.6 *±* 17.4 *µ*M for PGA2 and 21.9*±*2.6 *µ*M for 20*α*-dihydrocortisol; cortisol binding was not detectable (mean*±*s.d.; *n* = 3 independently prepared protein samples). **k–m**, Chemical structures and corrected candidate poses of PGA2 (**k**), 20*α*-dihydrocortisol (**l**) and cortisol (**m**) at NR4A3 binding site 1. **n**, NR4A3 model locating binding site 1; panels **k–n** all refer to this most stable candidate site. **o**, Generalized residue–ligand correlations at site 1 from triplicate atomistic simulations; circle colour and size encode correlation strength from 0.1 (yellow) to 1.0 (purple), with ligand atoms grouped below.

*Nr4a3*, encoding an orphan nuclear receptor implicated in context-dependent control of proliferation, differentiation and inflammation, was among the genes induced by 20*α*-dihydrocortisol [27, 28]. Knockdown of *Nr4a3* prevented induction of AP-1 and reduced *Myc*, *Bcl3*, *Egr2* and *Il6*; it also depleted NR4A3 protein and blocked the pERK/ERK response (Fig. 3d–g). Immunofluorescence independently confirmed loss of nuclear NR4A3 after knockdown (Extended Data Fig. 9). NR4A3 is therefore required for the measured epithelial response to the microbial steroid.

We next purified the mouse NR4A3 ligand-binding domain (Extended Data Fig. 10) and directly measured ligand binding. Prostaglandin A2 (PGA2), a reported NR4A3 ligand, bound with *K*_D_ = 66.6 *±* 17.4 *µ*M [29]. 20*α*-dihydrocortisol bound approximately threefold more tightly (*K*_D_ = 21.9 *±* 2.6 *µ*M), whereas cortisol showed no detectable binding (Fig. 3h–j). The cellular response observed at concentrations below the *K*_D_ measured for the isolated ligand-binding domain is consistent with signalling in an amplified intact-cell system, where full-length NR4A3, cofactors, local ligand partitioning or metabolism may influence effective receptor engagement. Docking and triplicate atomistic simulations identified binding site 1 as the most stable candidate pocket (Fig. 3k–o); the high-temperature retention test for this site is shown in Extended Data Fig. 11, and the complete alternative binding-site-2 analysis is shown in Extended Data Fig. 12. Generalized correlation analysis at site 1 further identified a ligand-specific interaction network centred on residues including Asn46 and Gln112 (Fig. 3o). Taken together, receptor loss of function, direct binding and atomistic modelling establish a change in molecular identity: reduction of one cortisol carbonyl generates an NR4A3-binding molecule with a host response that cortisol itself does not elicit.

### Human observations converge on the microbial pathway

We finally asked whether the pathway components co-occur in humans. The study included 134 adults identifying as Black or non-Hispanic White who underwent screening or surveillance colonoscopy in Chicago; subsets provided stool and hair for metagenomic, qPCR, steroid and cortisol analyses (Extended Data Fig. 13; Supplementary Tables 3–6). The self-identified groups differed in social and clinical variables, and Black participants were more likely to reside in neighbourhoods with greater poverty and homicide exposure (Fig. 4a,b) [30]. The group labels are therefore used to describe unequal lived exposures and are not interpreted as biological categories.

**Fig. 4.**
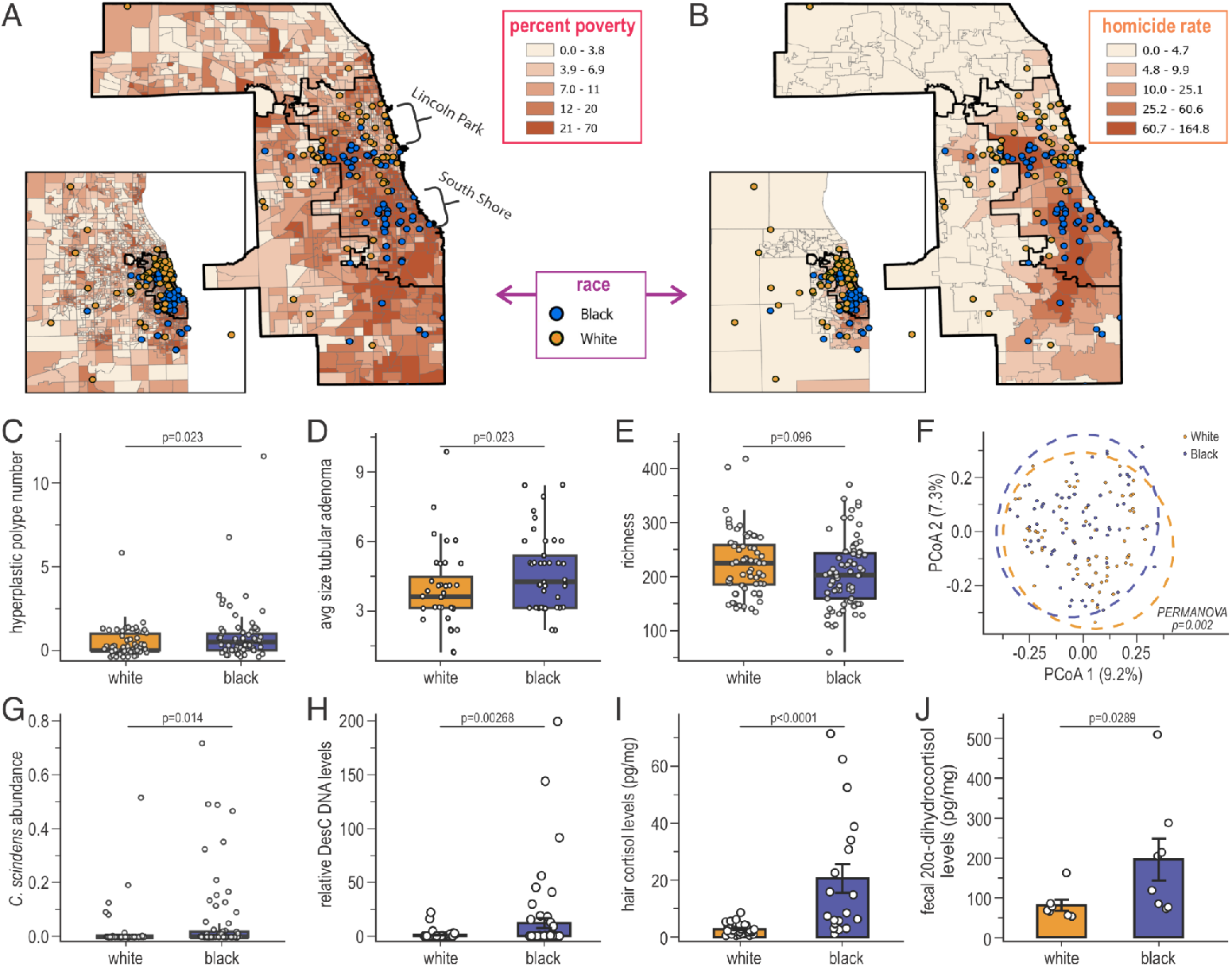
Cortisol exposure, the microbial DesC pathway and colorectal phenotypes converge in humans. **a,b**, Geocoded participant residences over census-tract percentage poverty (**a**) and community-area homicide rate per 100,000 residents (**b**), averaged from 2000–2018. Blue points denote participants self-identifying as Black and orange points denote non-Hispanic White participants; insets show the city-wide distribution, and bold boundaries indicate community areas, including South Shore and Lincoln Park. **c,d**, Hyperplastic polyp number (**c**) and average tubular adenoma size (**d**) among participants with the corresponding colonoscopy measure. **e**, Species richness in faecal metagenomes from White (orange; *n* = 63) and Black (blue; *n* = 71) participants. **f**, Principal-coordinate analysis of Bray–Curtis dissimilarity for the same metagenomes; points and dashed ellipses are orange for White and blue for Black participants, and axes report explained variance. Community composition was compared by PERMANOVA with 999 permutations. **g**, Relative abundance of *Clostridium scindens* in the metagenomic cohort (White, *n* = 63; Black, *n* = 71). **h**, Relative faecal *desC* DNA measured by qPCR (White, *n* = 53; Black, *n* = 58). **i**, Hair cortisol concentration (White, *n* = 28; Black, *n* = 19). **j**, Faecal 20*α*-dihydrocortisol among samples above the detection limit (White, *n* = 7; Black, *n* = 8); bars show mean*±*s.e.m. and points are participants. In **c–e,g–i**, each point is one participant; orange and purple denote White and Black participants, respectively. Boxes show medians and interquartile ranges, with whiskers extending to 1.5 times the interquartile range. Except for **f**, indicated *P* values are from two-sided Wilcoxon rank-sum tests. Self-identified group labels represent unequal lived and environmental exposures and are not biological categories.

At the population level, Black participants had more hyperplastic polyps and larger tubular adenomas in unadjusted comparisons (both *P* = 0.023; Fig. 4c,d). Faecal species richness showed a modest difference (*P* = 0.096), whereas overall community composition separated significantly (Bray–Curtis PERMANOVA *P* = 0.002; Fig. 4e,f). Within this altered microbiome, both *C. scindens* abundance (*P* = 0.014) and faecal *desC* DNA (*P* = 0.00268) were higher (Fig. 4g,h). Hair cortisol, which integrates longer-term hypothalamic–pituitary–adrenal-axis activity, was also markedly higher (*P <* 0.0001; Fig. 4i). These measurements place sustained glucocorticoid exposure and enrichment of the microbial cortisol-reduction pathway in the same human population.

Among samples with detectable metabolite, faecal 20*α*-dihydrocortisol was likewise higher (*P* = 0.0289; Fig. 4j). Thus, chronic cortisol exposure, the microbial organism and gene, its product and colorectal phenotypes are all represented in the human cohort. Together with the atomic, gnotobiotic and receptor-level experiments, these observations extend microbial cortisol reduction into a clinically relevant human setting.

## Discussion

This work shows that microbiome-dependent steroid metabolism can create signalling capacity rather than simply remove it. DesC converts cortisol—a glucocorticoid with no detectable binding to the NR4A3 ligand-binding domain—into 20*α*-dihydrocortisol, which binds NR4A3, activates an NR4A3-dependent ERK programme and changes intestinal transcription in vivo. The causal core is supported at four levels: an experimentally validated atomic mechanism; an isogenic catalytic-null bacterial control; metabolite production in the gut and circulation; and receptor loss of function coupled to direct binding.

The result expands the concept of the microbiome as an endocrine organ. Microbial metabolism of bile acids and other steroids is already known to diversify host ligands [2, 3, 6–8]; the DesC pathway provides a complete example in which a single stereospecific microbial reaction changes receptor selectivity. It also establishes a general strategy for assigning function to long-known but physiologically orphan microbial metabolites: resolve the enzyme mechanism, use that mechanism to build a catalytic control, follow the product in a defined host and identify the host receptor.

The colorectal observations place this mechanism in a disease-relevant human context. Although the cohort does not establish a causal pathway or explain health disparities, the convergence of cortisol exposure, faecal *desC*, 20*α*-dihydrocortisol and colorectal phenotypes identifies a tractable hypothesis for prospective testing in chronic stress, Cushing syndrome and therapeutic glucocorticoid exposure, settings independently associated with increased cancer risk [31–33]. More broadly, our findings establish that microbial steroid metabolism can generate biologically distinct host signals and raise the possibility that many products currently catalogued as terminal microbial metabolites may instead represent unrecognized components of host–microbiome signalling.

## Methods

### Protein production and biochemical assays

The open reading frame encoding *C. scindens* ATCC 35704 DesC (EDS07887) was cloned into pET-51b and expressed in *E. coli* BL21 CodonPlus (DE3)-RIPL as described previously [9]. Site-directed variants were generated with the primers in Supplementary Table 7, confirmed by Sanger sequencing, and purified by the same Strep-Tactin procedure as wild-type DesC. Human 20*α*-hydroxysteroid dehydrogenase (20*α*-HSDH; AKR1C1) and the mouse NR4A3 ligand-binding domain were expressed as N-terminal His-tagged proteins in *E. coli* BL21 and purified by nickel-affinity chromatography. Protein purity and apparent molecular mass were assessed by SDS–PAGE.

DesC activity was measured from NADH oxidation at 340 nm (*ε* = 6, 220 M*^−^*^1^cm*^−^*^1^) in 50 mM phosphate, pH 7.5, containing cortisol, NADH and enzyme. Initial rates were fitted by nonlinear regression to the Michaelis–Menten equation. Substrate profiling used 200 *µ*M steroid and three biological replicates per substrate. AKR1C1 activity was measured analogously from NADPH oxidation in 100 mM potassium phosphate, pH 7.0, and normalized to progesterone. Circular-dichroism spectra of DesC variants were collected at 25 °C from 190 to 260 nm using 0.2 mg ml*^−^*^1^ protein in 10 mM KH_2_PO_4_, pH 7.5, with five accumulations.

DesC binding was measured on a MicroCal VP-ITC instrument. Proteins were dialysed against 50 mM sodium phosphate, 150 mM NaCl and 10% glycerol, pH 7.5. Protein (50 *µ*M), with or without NADH, was titrated with cortisol and fitted to a single-site model. NR4A3 binding was measured on a PEAQ-ITC instrument in 50 mM sodium phosphate, 150 mM NaCl and methyl-*β*-cyclodextrin, pH 7.5. NR4A3 ligand-binding domain (100 *µ*M) was titrated with 13 successive 3-*µ*l injections of 2 mM PGA2, 20*α*-dihydrocortisol or cortisol at 150-s intervals. Three independently prepared protein samples were analysed for each ligand.

### DesC crystallography

Selenomethionine-labelled DesC was crystallized by sitting-drop vapour diffusion at 20 °C against 0.2 M MgCl_2_, 24% PEG 400 and 0.1 M HEPES, pH 6.75. Crystals were cryoprotected with 10% 1,2-ethanediol and flash frozen. Diffraction data were collected at beamline BL14-1 of the Stanford Synchrotron Radiation Lightsource, indexed with XDS, scaled with XSCALE, phased by selenium multiwavelength anomalous dispersion with SHELX, and refined with Refmac [34–36]. Data-collection and refinement statistics are in Supplementary Table 2. Coordinates are deposited under PDB accession 4OH1.

### DesC molecular dynamics and QM/MM calculations

NADH was positioned by structural alignment to homologous cofactor-bound medium-chain dehydrogenases, and cortisol was placed with its C20 carbonyl adjacent to the NADH hydride donor and catalytic zinc. Five independent models were prepared with VMD and QwikMD [17, 37], solvated with explicit TIP3P water and 0.15 M NaCl, and simulated with NAMD under periodic NpT conditions at 300 K and 1 bar [18]. CHARMM36 parameters, particle-mesh Ewald electrostatics, a 12-Å short-range cutoff and a 2-fs integration step were used. Each of the five independent replicas underwent minimization and 10 ns of restrained relaxation followed by 100 ns of unrestrained MD, providing a total of 500 ns of unrestrained sampling. All five replicas converged to a similar stable DesC–NADH–cortisol active-site configuration.

A representative configuration from the conformational state consistently sampled across the five classical MD replicas was used to initiate electrostatically embedded QM/MM simulations through the NAMD interface [20]. The MM region used CHARMM36; the 573-atom QM region used PM7 and contained the substrates, catalytic zinc and surrounding active-site residues. Thirty-six QM–MM boundaries were treated with link atoms. After unbiased QM/MM equilibration, candidate pathways were generated by QM/MM steered MD that biased only the initiating bond changes.

The selected pathway was subjected to extensive ensemble-based enhanced sampling. Finite-temperature string optimization used 25 images and 750 independently propagated QM/MM replicas per string iteration, distributed along the reaction path. Collective variables described the donor–hydrogen and acceptor–hydrogen distances associated with hydride transfer and the three proton transfers. Free energy along the optimized pathway was then estimated by multiple-walker extended adaptive biasing force calculations using 600 walkers, with 10,000 MD steps per walker and path variables *S* and *Z* [21–23]. Thus, the reaction pathway and free-energy profile derive from ensemble-based sampling across hundreds of QM/MM trajectories rather than from a single transition trajectory. Reported free energies describe the modelled chemical transformation and do not include substrate binding, conformational closure or product release.

### NR4A3 structural modelling

The full-length mouse NR4A3 model was generated with AlphaFold 3. PGA2, 20*α*-dihydrocortisol and cortisol were protonated at pH 7.4 and independently docked into two candidate binding pockets. Each ligand–pocket complex was solvated in explicit TIP3P water with 0.15 M NaCl and simulated in three independent replicas with NAMD 3. Following minimization, each replica was gradually heated to 300 K, equi-librated for 0.5 ns and subjected to production MD at 300 K. Thus, ligand stability and binding-site behaviour were assessed across three independent trajectories for each ligand–pocket combination. Additional temperature-ramp simulations were also performed in triplicate and were used exclusively as qualitative stress tests of ligand retention. Ligand–pocket distances and generalized residue–ligand correlations were analysed across the three independent replicas. These simulations were used to assess the reproducibility and stability of the predicted binding modes and to generate structural hypotheses; they were not used to estimate binding affinities.

### Construction and validation of engineered bacteria

Wild-type *desC* and the S47A allele were integrated into the SS3 intergenic locus between *ompW* and *yciE* in *E. coli* BL21(DE3). A pCas/pTarget dual-plasmid CRISPR–Cas9 system supplied Cas9, *λ*-Red recombination and the locus-specific guide RNA [38]. Donor cassettes contained the J23100 constitutive promoter, pET ribosome-binding site, His tag, *desC* allele and T7 terminator flanked by 50-bp homology arms. Integration was confirmed by PCR and sequencing, and both editing plasmids were cured by counterselection and nonselective growth. An EGFP cassette was used initially to validate stable expression from the locus.

Eco_desCWT and Eco_desCM were cultured from equal starting density in LB containing 50 *µ*M cortisol and 5 g l*^−^*^1^ glycerol. Growth and glycerol utilization were monitored, and cortisol and 20*α*-dihydrocortisol were quantified by reverse-phase HPLC with diode-array detection at 254 nm. Plasmids, primers and donor sequences are listed in Supplementary Tables 8–10.

### Gnotobiotic mouse experiment

All animal procedures were approved by the University of Illinois Urbana–Champaign IACUC (protocol 19222). Germ-free C57BL/6J mice were maintained in flexible-film isolators and screened routinely for contamination. Individually caged mice were gavaged with 10^8^ colony-forming units of Eco_desCWT or Eco_desCM on days 1 and 3. From days 4–7, mice received sterile water with or without cortisol (10 mg kg*^−^*^1^ day*^−^*^1^); four mice were analysed per strain-by-treatment group. Colonization and mono-association were confirmed from caecal material by V4 16S rRNA gene sequencing and QIIME 2 analysis [39]. On day 7, mice were euthanized and serum, caecal contents and intestinal epithelial scrapings were collected.

Steroids were extracted from serum and caecal homogenates by supported liquid extraction with methyl-*tert*-butyl ether. Reconstituted samples were separated by two-dimensional liquid chromatography and quantified by positive-ion dynamic multiple-reaction monitoring on an Agilent 6495A triple-quadrupole mass spectrometer. Transitions and internal standards are listed in Supplementary Tables 1 and 2.

### Primary intestinal epithelial cells

Primary epithelial cells were isolated from conventionally raised C57BL/6 mouse intestine by collagenase and hyaluronidase digestion, differential centrifugation and short-term culture in complete DMEM. Before steroid exposure, medium was replaced with charcoal-stripped serum. Cells were treated with vehicle, cortisol or 20*α*-dihydrocortisol (500 nM) for 20 min or 6 h. For loss-of-function experiments, cells at 40–50% confluence were transfected with 40 pmol per well of *Nr4a3* siRNA or control siRNA using Lipofectamine 2000, incubated for 48 h and then exposed to vehicle or 20*α*-dihydrocortisol for 6 h.

RNA was isolated with TRI Reagent, DNase treated and purified. cDNA was quantified by SYBR Green qPCR relative to *β*-actin using the 2*^−^*^Δ*Cq*^ method; primer sequences are in Supplementary Table 13. For immunoblotting, 50 *µ*g protein per lane was separated on 12% SDS–PAGE, transferred to nitrocellulose and probed for NR4A3, ERK, pERK and *β*-actin. Bands were detected by chemiluminescence and quantified with iBright software. For immunofluorescence, fixed and permeabilized cells were stained for CDX2, pERK or NR4A3, followed by Alexa Fluor 488 secondary antibody and Hoechst nuclear stain. Nuclear localization was quantified as the percentage of marker-positive nuclei.

### RNA sequencing and pathway analysis

RNA from mouse intestinal scrapings and primary epithelial cells was quality assessed and converted to indexed poly(A)-selected libraries. Mouse-tissue libraries were sequenced as paired-end 150-nt reads and epithelial-cell libraries as single-end 100-nt reads on an Illumina NovaSeq 6000. Adaptors and low-quality sequence were removed with Trimmomatic, residual rRNA was filtered with SortMeRNA, and transcript abundance was quantified with Salmon [40–42]. Counts were filtered and normalized with edgeR, differential expression was tested with limma, and KEGG enrichment was performed with clusterProfiler [43–45]. False-discovery-rate correction and contrasts are specified in the figure legends and Source Data.

### Human cohort

Adults aged 45–75 years who self-identified as Black or White were recruited during screening or surveillance colonoscopy at Rush University Medical Center and the University of Illinois Chicago. Exclusion criteria included current or recent malignancy, previous colorectal cancer, major gastrointestinal disease, recent antibiotics, regular prebiotic or probiotic supplementation, conditions or medications affecting digestion, pregnancy or nursing, and recent menstruation. The study was approved by the institutional review boards of Rush University Medical Center (13102201), UIC/UIUC (2016-0495) and Purdue University (2022-1626); all participants provided written informed consent. Of 161 participants completing the study, 134 had metagenomic data; assay-specific sample sizes and missingness are reported in Supplementary Tables 3–6.

Hair was sampled from the crown vertex approximately 30 days after colonoscopy. The proximal 3 cm was washed, ground, extracted in methanol and assayed using a DetectX cortisol enzyme immunoassay following established quality controls [46]. Implausible values, values more than three standard deviations from the mean and samples from participants reporting corticosteroid-product use were excluded according to the prespecified rules.

Stool collected approximately four weeks after colonoscopy was stored at *−*80 *^◦^*C. DNA was extracted with the DNeasy PowerSoil kit and sequenced as paired-end 150-bp shotgun libraries after quality filtering and human-read depletion. Species profiles were generated with MetaPhlAn 4 [47]. Faecal *desC* was quantified by SYBR Green qPCR relative to universal 16S rRNA gene primers using 2*^−^*^Δ*Cq*^; primers are listed in Supplementary Table 15. Lyophilized stool was extracted in methanol, spiked with internal standards and analysed for 20*α*-dihydrocortisol by the LC–MS/MS workflow used for mouse samples.

Participant addresses were geocoded and linked to community-level poverty and homicide estimates from the Chicago Health Atlas. Self-identified group was used as a social exposure descriptor and not as a biological variable. The cohort is observational; analyses did not test a causal effect of race.

### Statistics

Statistical analyses were performed in R 4.4.3 or GraphPad Prism as specified. Data distributions were assessed with the Shapiro–Wilk test. Two-group continuous comparisons used two-sided Wilcoxon rank-sum tests when distributional assumptions were not met; gene and protein measurements used two-sided *t*-tests with Holm correction. Pathway and other families of tests used Benjamini–Hochberg false-discovery-rate correction. Species richness was compared by Wilcoxon rank-sum test. Bray–Curtis community dissimilarity was tested by PERMANOVA with 999 permutations. Because many human stool samples were below the 20*α*-dihydrocortisol detection limit, detection frequency was first compared by Fisher’s exact test and concentrations were then compared only among detectable samples by Wilcoxon rank-sum test. Exact *n*, statistical tests, multiplicity corrections and definitions of centre and error bars are given in the corresponding legends and Source Data.

## Acknowledgements

J.M.R. would like to express gratitude to both the Cancer Center at Illinois and the Center for Advanced Study at Illinois for the financial support and protected time to pursue this work. We want to thank Alvaro Hernandez, Director of the DNA Services Facility at the Roy J. Carver Biotechnology Center at Urbana-Champaign, and Christopher J. Fields, Director of High-Performance Biology Computing at the Roy J. Carver Biotechnology Center, for nucleic acid sequencing and bioinformatics assistance. We thank Furong Sun, Director of Mass Spectrometry Laboratory at Urbana-Champaign for LC-MS analysis. All chemical structures were created with ChemDoodle. This work was supported by grants from the National Institutes of Health R01GM145920 [J.M.R, H.R.G], R01GM134423 [J.M.R., A.J.S., Y.J.], R01CA204808 [H.R.G., J.M.R., E.M., L.T-H], U54MD012523 [L.T-H., J.M.R., H.R.G], and R01GM086596 [R.J.A.]. J.M.R. was also supported by UIUC Department of Animal Sciences Matchstick grant and Hatch ILLU-538-916. B.B. was supported by NIGMS Diversity Supplement on R01 GM134423. Computational work supported by the National Science Foundation grant MCB-2143787 and NIH R24 GM145965 [R.C.B.]. The authors acknowledge financial support from the Alabama Graduate Research Scholars Program (GRSP) funded through the Alabama Commission for Higher Education and administered by the Alabama EPSCoR. We thank the members of the JCSG high-throughput structural biology pipeline for their contribution to this work. The structural determination work (PDB ID: 4OH1) was supported by the NIH, National Institute of General Medical Sciences (NIGMS), Protein Structure Initiative [U54 GM094586]. Portions of this research were carried out at the Stanford Synchrotron Radiation Lightsource, SLAC National Accelerator Laboratory, supported by the U.S. Department of Energy, Office of Science, Office of Basic Energy Sciences under Contract No. DE-AC02-76SF00515. The SSRL Structural Molecular Biology Program is supported by the DOE Office of Biological and Environmental Research, and by the National Institutes of Health, National Institute of General Medical Sciences (P41GM103393). The contents of this publication are solely the responsibility of the authors and do not necessarily represent the official views of NIGMS or NIH.

## Author contributions

J.M.R. conceptualized and designed the overall research with important contributions to the clinical studies by L.T-H., and P. G. W., bacterial engineering from J.L., and molecular dynamics by R.C.B. M.F., T.W., R.S.R., A.O., M.C.R.M., D.S., P.S.B., L.Y., A.M.A-H., H.L.D., S.D., R.M.P., S.G., L.R., A.H., N.K., S.M.M., S.B., S.A.L., P.B.H., Z.L-S., F.P., E.M., H.R.G., Y-S.J., S.J.K., P.G., R.J.A., I.C., A.J.S., M.R., N.M.K., and S.O., performed the experiments and analyzed the data. J.M.R., L.T-H., P.G.W., and J.W.L., supervised the study. J.M.R., M.F., T.W., R.C.B., L.T-H., and P.G.W., wrote the manuscript with input from all authors. M.F. and T.W. contributed equally to this work. All authors edited the manuscript and approved the final draft.

## Data availability

The DesC crystallographic coordinates and associated structure factors are available from the Protein Data Bank under accession code 4OH1. The 16S rRNA gene V4 amplicon sequencing data generated in this study have been deposited in the NCBI Sequence Read Archive under BioProject accession PRJNA1469766. The RNA-sequencing data have been deposited under BioProject accession PRJNA1470094.

During peer review, the deposited sequencing datasets can be accessed through the following private NCBI reviewer links:

- 16S rRNA gene V4 amplicon sequencing (PRJNA1469766): https://dataview.ncbi.nlm.nih.gov/object/PRJNA1469766?reviewer=2kqt2nvukndg98sfibdr7vpq1b
- RNA sequencing (PRJNA1470094):

https://dataview.ncbi.nlm.nih.gov/object/PRJNA1470094?reviewer=1b8dpg6api7da9s9fhh0hegg6f

Source data underlying the quantitative graphs, together with uncropped immunoblots and other primary data required to reproduce the reported analyses, are provided with this paper as Source Data. Additional experimental results, computational analyses and supporting data are provided in the Supplementary Information and Extended Data. All other data supporting the findings of this study are available within the Article and its Supplementary Information or from the corresponding authors upon reasonable request.

## Code availability

No custom code was developed for this study. All computational analyses were performed using established software packages as described in the Methods. The input structures, simulation configuration files, parameters and collective-variable definitions, together with the files required to reproduce the DesC and NR4A3 computational analyses, are provided with this paper as Supplementary Data 1 (ZIP archive).

## Additional information

Supplementary Information is available for this paper. Correspondence and requests for materials should be addressed to Jason M. Ridlon.

## Extended Data

**Figure ED1:**
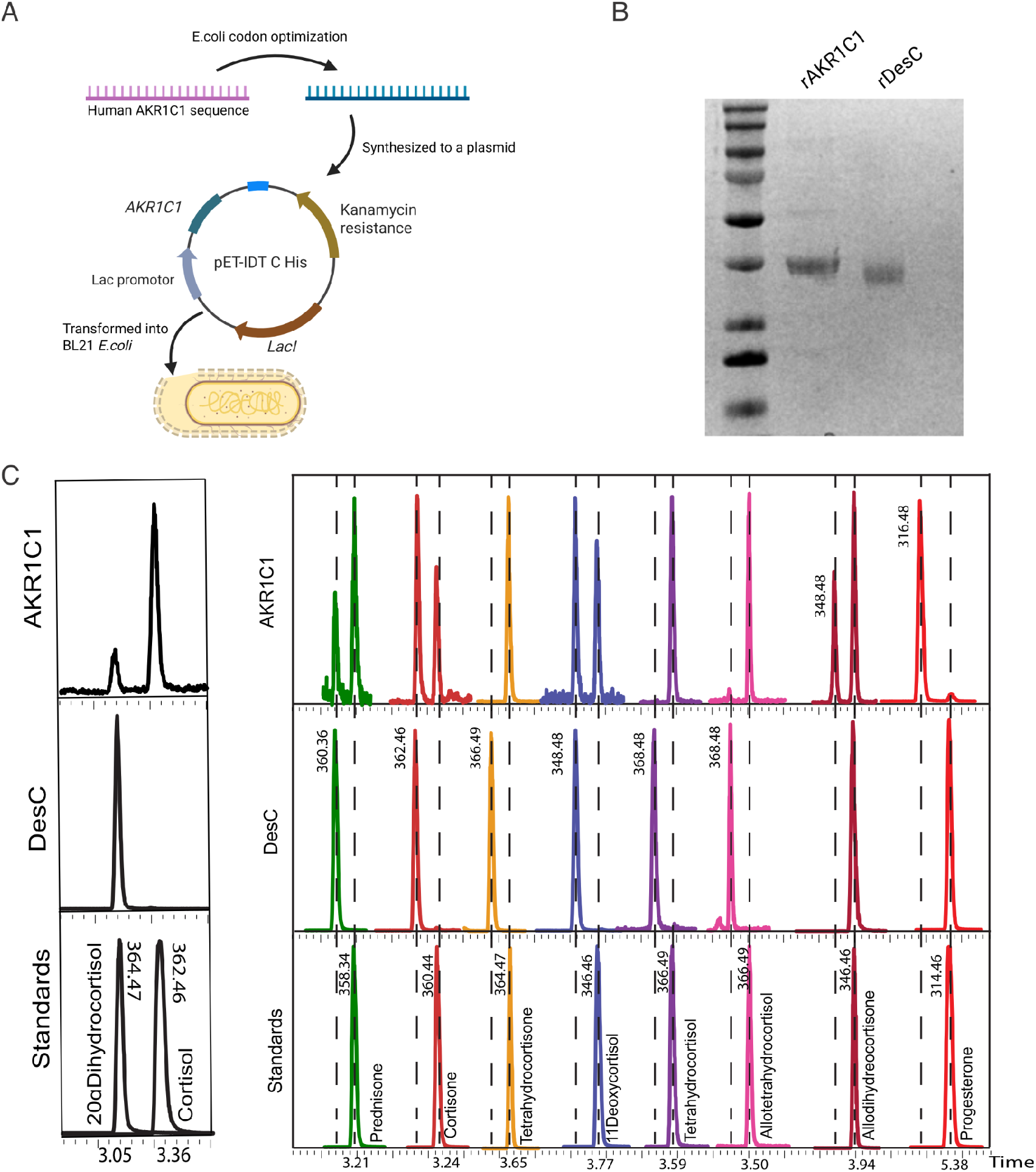
Recombinant human AKR1C1 production and chromatographic validation of steroid products. **a**, Human *AKR1C1* was codon-optimized for *E. coli*, synthesized in a pET-IDT C-His expression vector and transformed into BL21 cells for recombinant expression. The schematic marks the *AKR1C1* insert, lac regulatory elements and kanamycin-resistance cassette. **b**, Representative SDS–PAGE of purified recombinant human AKR1C1 (rAKR1C1) and recombinant bacterial DesC (rDesC); the molecular-mass ladder is shown at left. **c**, Representative chromatographic traces used to establish steroid retention times and assign products in the DesC and AKR1C1 substrate-specificity assays. Experimental enzyme traces are shown above the corresponding authentic steroid standards; coloured traces denote the individual steroids and dashed vertical markers denote the assigned retention positions. These controls support the distinct substrate profiles summarized in Fig. 1b.

**Figure ED2:**
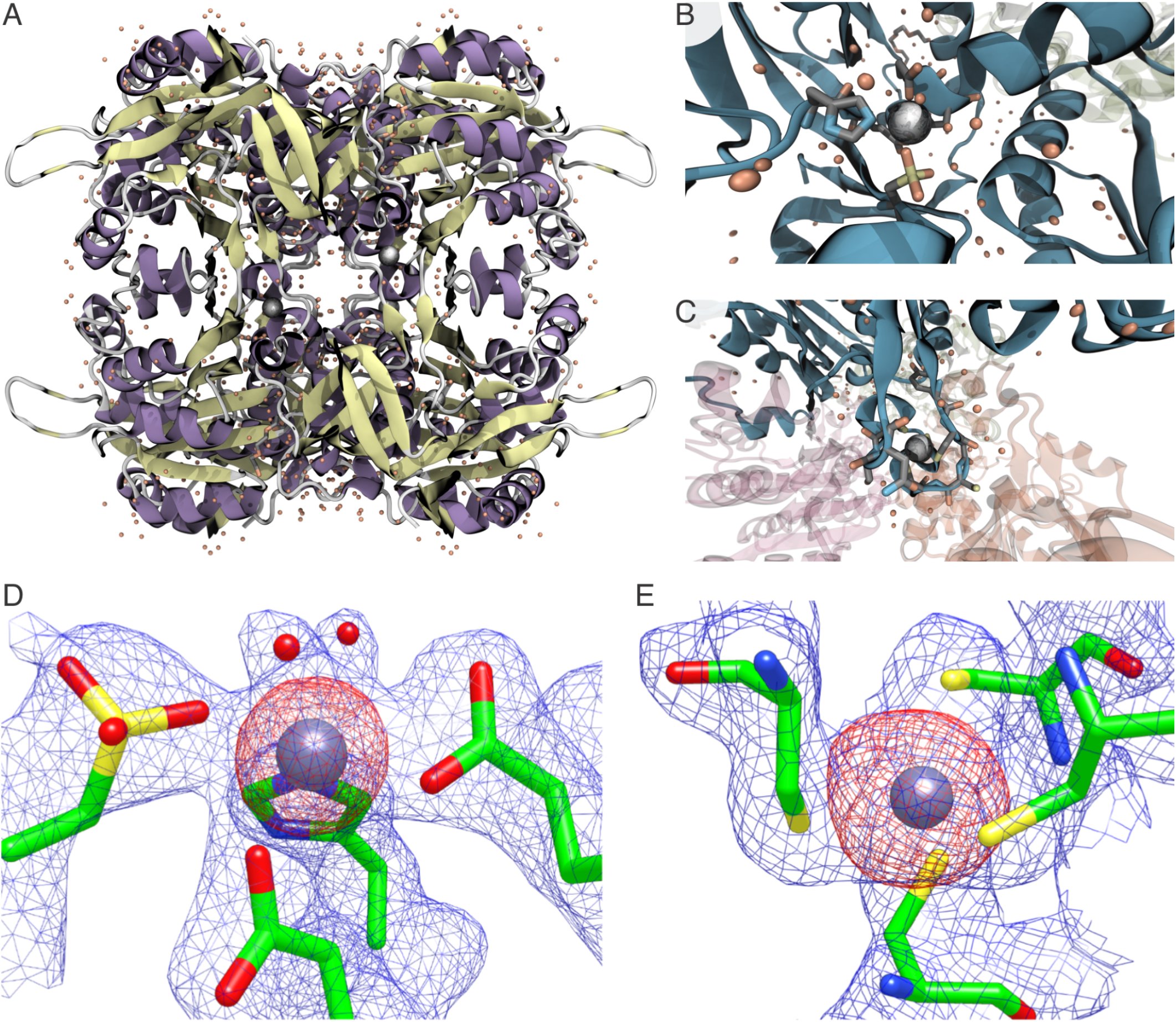
DesC quaternary structure and electron density at the two zinc sites. **a**, Tetrameric DesC crystal structure viewed along the oligomeric axis; secondary-structure elements are coloured by type, crystallographic waters are shown as orange spheres and zinc ions as grey spheres. **b**, Close view of the catalytic zinc environment and surrounding solvent in one DesC monomer. **c**, Structural zinc at the extended lobe that contributes to the oligomeric interface; neighbouring monomers are shown translucently. **d**, Electron-density maps surrounding the catalytic zinc and its coordinating residues. The zinc *F_o_− F_c_* difference density is shown as red mesh contoured at 7*σ*, and the 2*F_o_ − F_c_* density for zinc and coordinating residues as blue mesh contoured at 1.3*σ*. **e**, Corresponding maps for the structural zinc site, with *F_o_ − F_c_* density contoured at 5*σ* and 2*F_o_ − F_c_* density at 1.0*σ*. Crystallographic data and refinement statistics are given in Supplementary Table 2a.

**Figure ED3:**
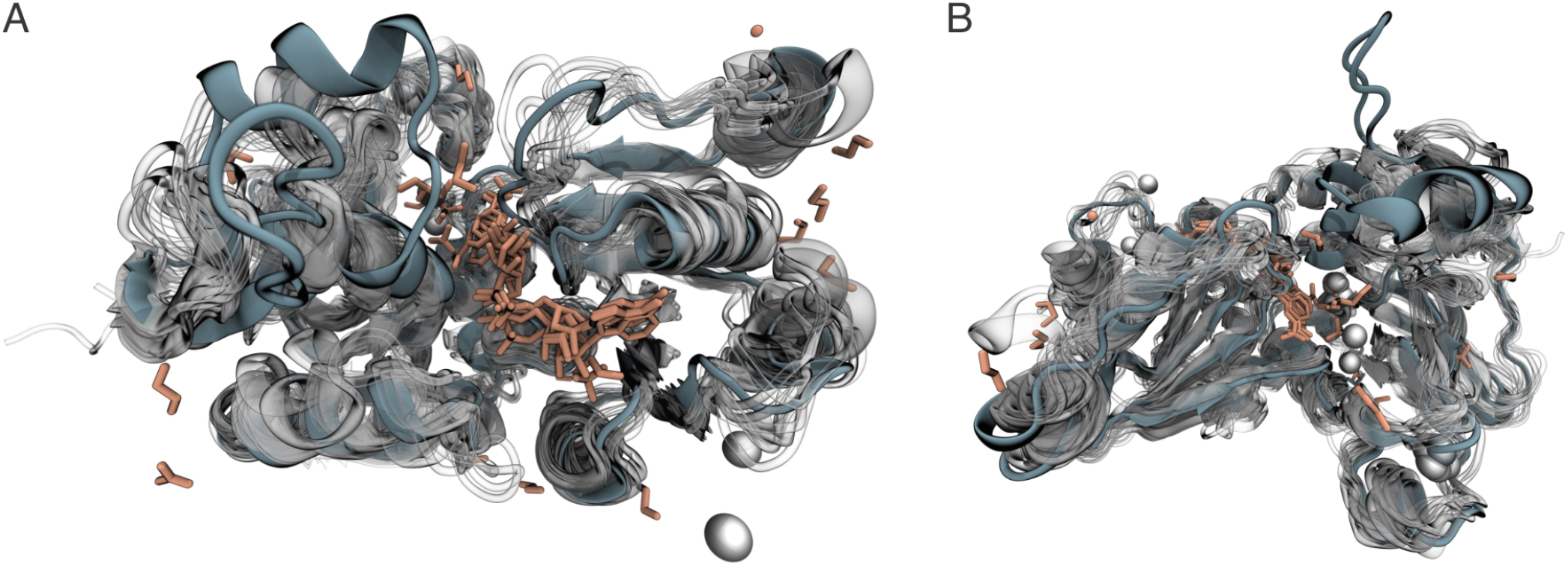
Structural alignment of DesC with cofactor-bound medium-chain dehydrogenase homologues. **a,b**, Superposition of the apo-DesC structure with structurally related, cofactor-bound dehydrogenases in two orthogonal orientations. DesC is highlighted in blue-grey; aligned homologues are shown as transparent grey cartoons, bound cofactors and ligands as sticks, solvent as orange spheres and zinc ions as spheres. The conserved cofactor geometry across the aligned structures was used to position NADH in DesC before cortisol placement and classical molecular-dynamics refinement.

**Figure ED4:**
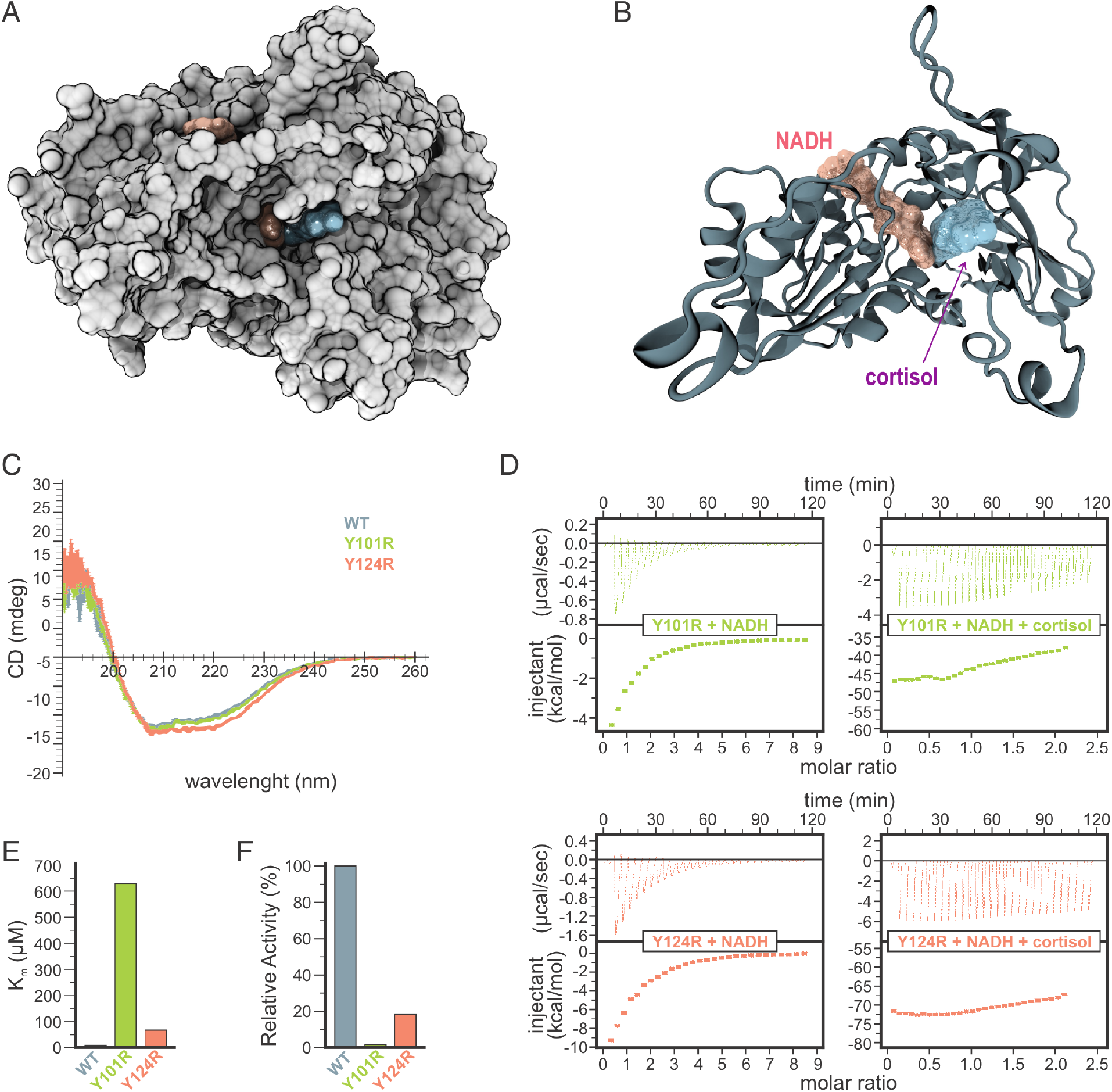
Experimental validation of the MD-predicted cortisol-recognition residues Tyr101 and Tyr124. **a**, Molecular surface of DesC showing the NADH/cortisol-containing active-site cleft. **b**, Cartoon representation of the same pocket with NADH and cortisol shown as surfaces, illustrating the spatial relationship of the cofactor and steroid. **c**, Circular-dichroism spectra of wild-type DesC (WT, blue-grey), Y101R (green) and Y124R (salmon), showing preservation of the global secondary-structure profile after mutation. **d**, Representative isothermal-titration-calorimetry experiments for Y101R (top) and Y124R (bottom) with NADH alone (left) or NADH plus cortisol (right). **e**, Apparent Michaelis constants for cortisol for WT, Y101R and Y124R DesC. **f**, Relative catalytic activity of the same proteins under saturating cofactor and substrate conditions. Y101R and Y124R lost 96.0*±*0.7% and 81.8*±*0.3% of WT activity, respectively. Kinetic values and secondary-structure estimates are reported in Supplementary Tables 2c,d.

**Figure ED5:**
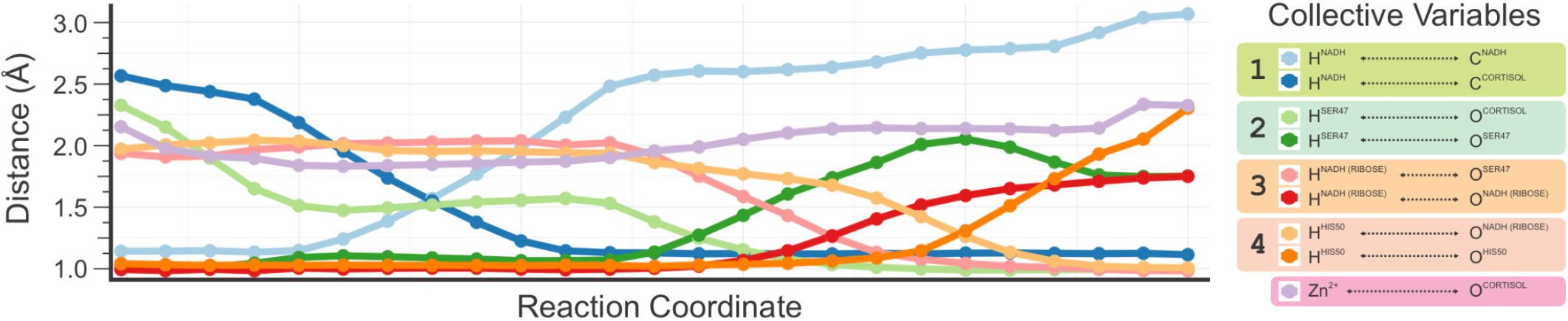
Collective variables describing the optimized DesC reaction coordinate. Interatomic distances for the collective variables used to follow the four bond rearrangements in the QM/MM string calculation are plotted across the reaction coordinate. Paired shades track donor–hydrogen and hydrogen–acceptor distances for hydride transfer from NADH to cortisol and the three subsequent proton transfers that restore the catalytic protonation state; the zinc–substrate distance is shown separately. The coordinated changes resolve hydride transfer, protonation of the cortisol carbonyl oxygen and the subsequent Ser47/ribose/His50 proton relay that underlies the pathway shown in Fig. 1f,i,j.

**Figure ED6:**
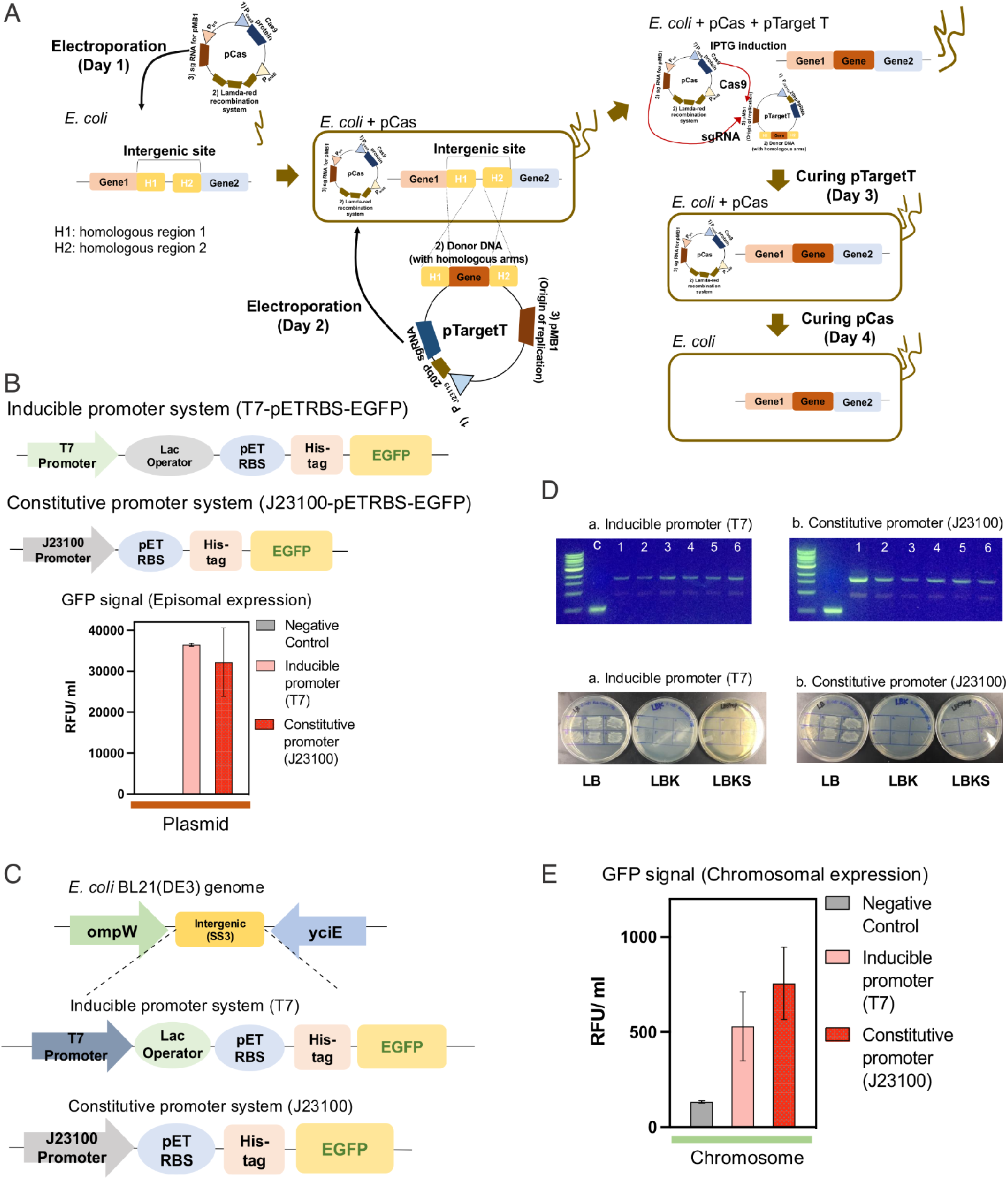
Validation of stable chromosomal expression in the engineered *E. coli* background. **a**, pCas/pTarget CRISPR–Cas9 workflow used to integrate donor DNA into an intergenic chromosomal locus by *λ*-Red-assisted homologous recombination, followed by sequential curing of pTarget and pCas. **b**, EGFP reporter cassettes used to compare inducible T7 and constitutive J23100 promoter architectures, with episomal reporter fluorescence shown below. **c**, SS3 intergenic locus between *ompW* and *yciE* and the corresponding inducible and constitutive reporter inserts used to validate chromosomal expression. **d**, PCR verification of chromosomal integration (top) and replica-plating controls used to verify loss of the editing plasmids (bottom). **e**, EGFP signal after chromosomal integration of the two reporter architectures. This validated constitutive SS3 expression strategy was then used to generate the isogenic Eco_*desC* WT and Eco_*desC* M strains in Fig. 2a.

**Figure ED7:**
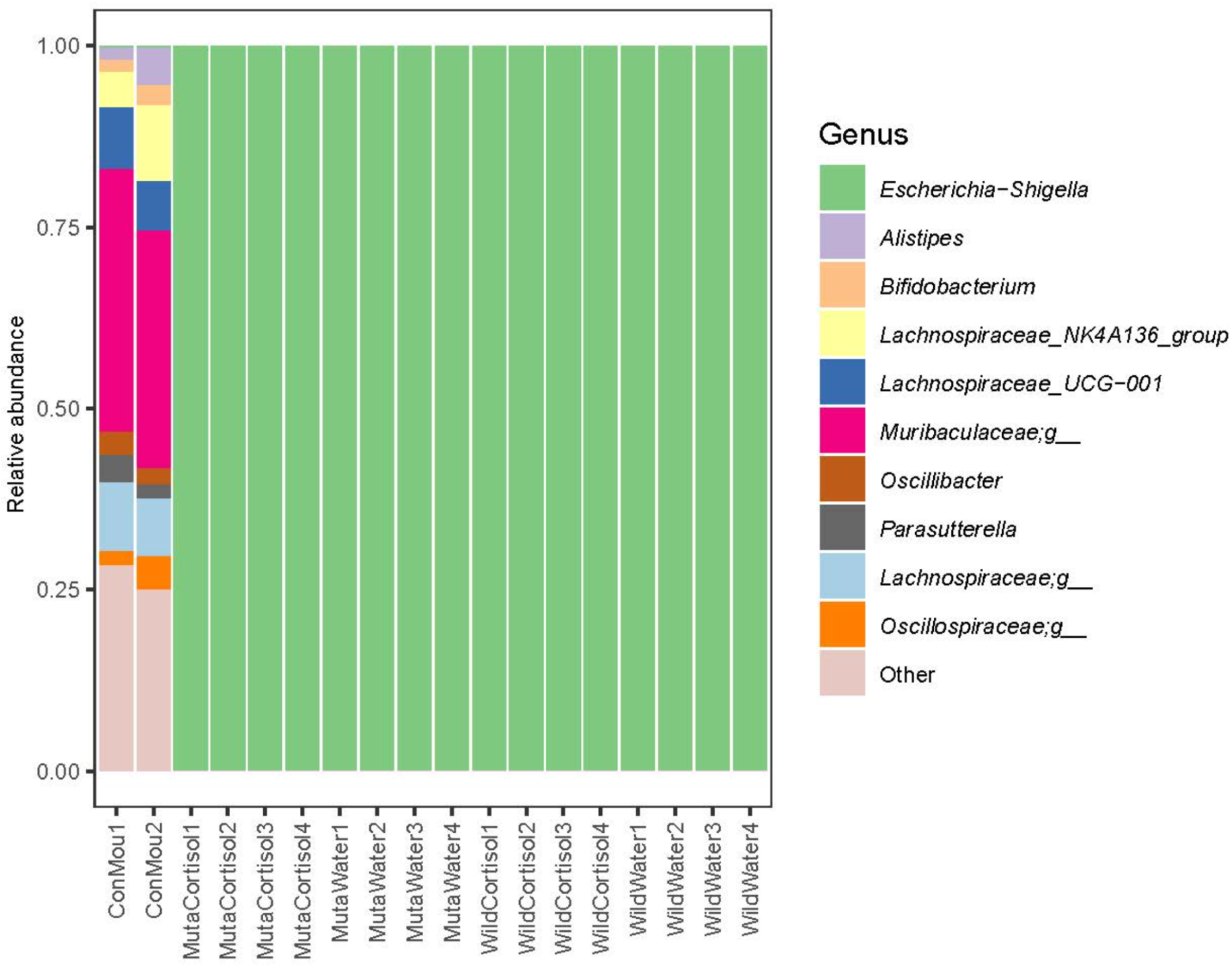
16S rRNA gene profiles confirm mono-association in the gnotobiotic experiment. Relative genus-level abundance from V4 16S rRNA gene sequencing of caecal DNA. The two comparison samples (ConMou1 and ConMou2) show mixed communities, whereas all experimental samples from mice colonized with engineered *E. coli* are dominated by *Escherichia–Shigella*. Experimental groups comprise mice colonized with Eco_*desC* M and given cortisol (MutaCortisol1–4) or water (MutaWater1–4), and mice colonized with Eco_*desC* WT and given cortisol (WildCortisol1–4) or water (WildWater1–4). Four analysed mice are shown per experimental group.

**Figure ED8:**
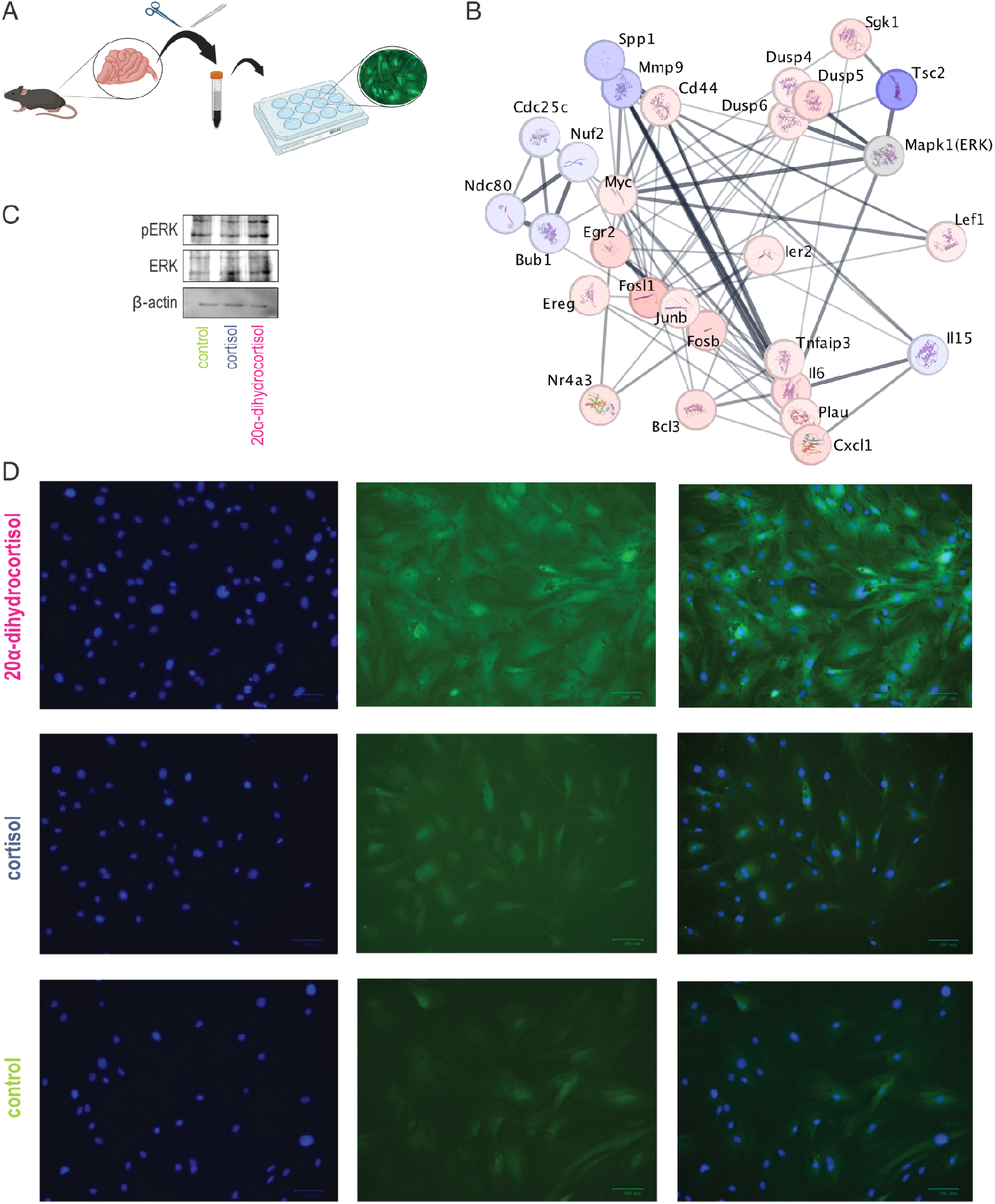
20*α*-dihydrocortisol induces an ERK-linked transcriptional programme in primary intestinal epithelial cells. **a**, Schematic of primary intestinal epithelial-cell isolation and culture. **b**, Functional interaction network among representative genes regulated after 6 h exposure to 500 nM 20*α*-dihydrocortisol; nodes mark regulated genes and edges denote functional associations. **c**, Representative immunoblots for pERK, total ERK and *β*-actin after vehicle, cortisol or 20*α*-dihydrocortisol treatment. **d**, Representative immunofluorescence fields after 20*α*-dihydrocortisol (top), cortisol (middle) or vehicle (bottom). Nuclei are blue, pERK is green and merged images are shown at right. Together with the quantitative measurements in Fig. 3b,c, these data show selective delayed ERK activation and nuclear translocation in response to the microbial steroid.

**Figure ED9:**
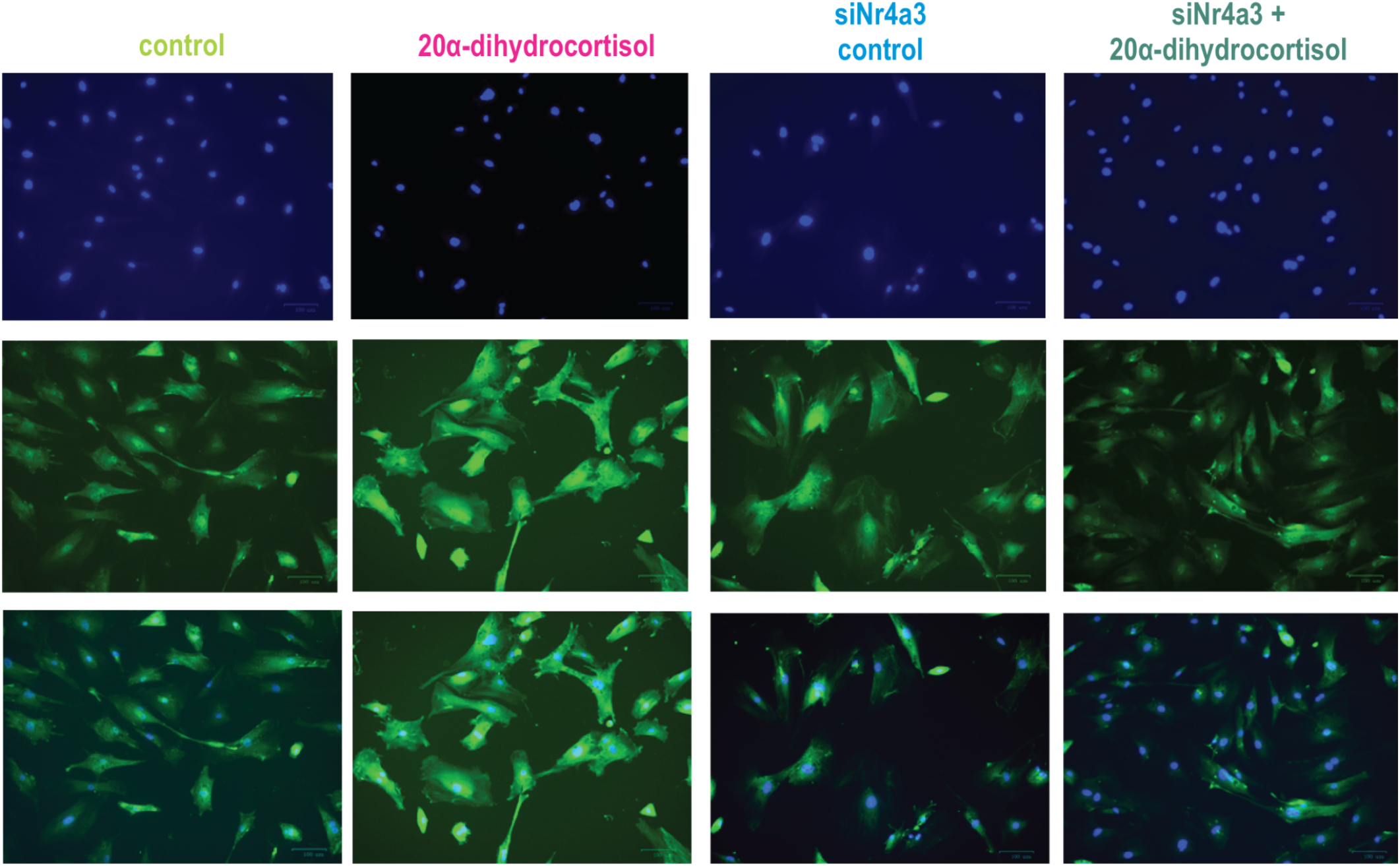
NR4A3 immunofluorescence after *Nr4a3* knockdown and 20*α*-dihydrocortisol exposure. Representative primary intestinal epithelial cells treated with control siRNA plus vehicle, control siRNA plus 20*α*-dihydrocortisol, *Nr4a3* siRNA plus vehicle or *Nr4a3* siRNA plus 20*α*-dihydrocortisol. Columns correspond to the four conditions indicated above the images. Nuclei are shown in blue (top row), NR4A3 immunoreactivity in green (middle row) and merged images in the bottom row. The loss of nuclear NR4A3 after siRNA treatment provides an imaging-level validation of the knockdown underlying the transcriptional and ERK loss-of-function experiments in Fig. 3d–g.

**Figure ED10:**
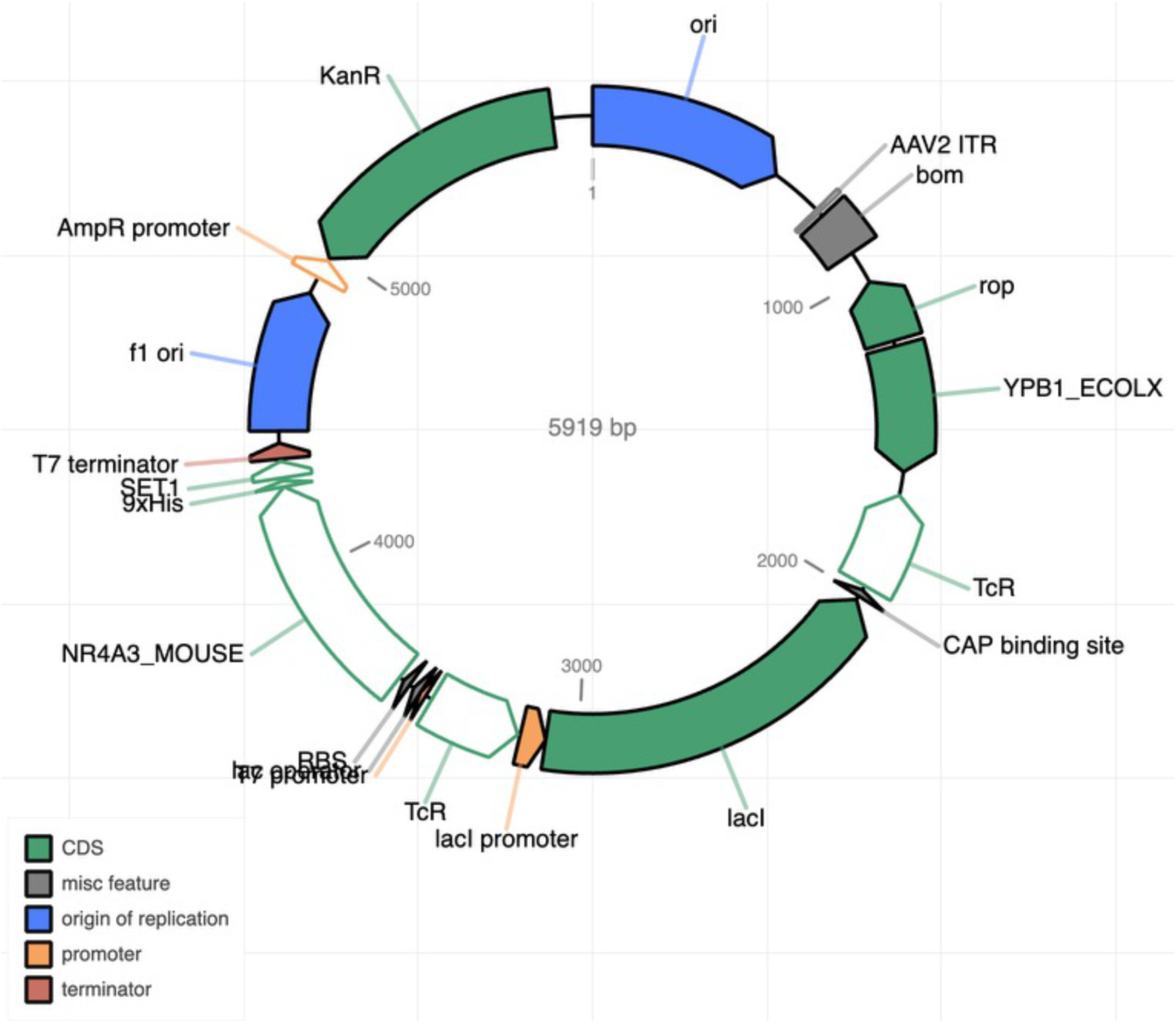
Expression construct for the mouse NR4A3 ligand-binding domain. Circular map of the 5,919-bp pET-IDT C-His expression plasmid used to produce the recombinant mouse NR4A3 ligand-binding domain for isothermal-titration calorimetry. The map shows the *NR4A3_MOUSE* insert, affinity tag, promoter/regulatory elements, replication origins and selectable-marker features. Recombinant NR4A3 produced from this construct was purified by nickel-affinity chromatography and used for the direct PGA2, 20*α*-dihydrocortisol and cortisol binding measurements in Fig. 3h–j. The amino-acid sequence of the expressed ligand-binding-domain construct is given in Supplementary Table 14.

**Figure ED11:**
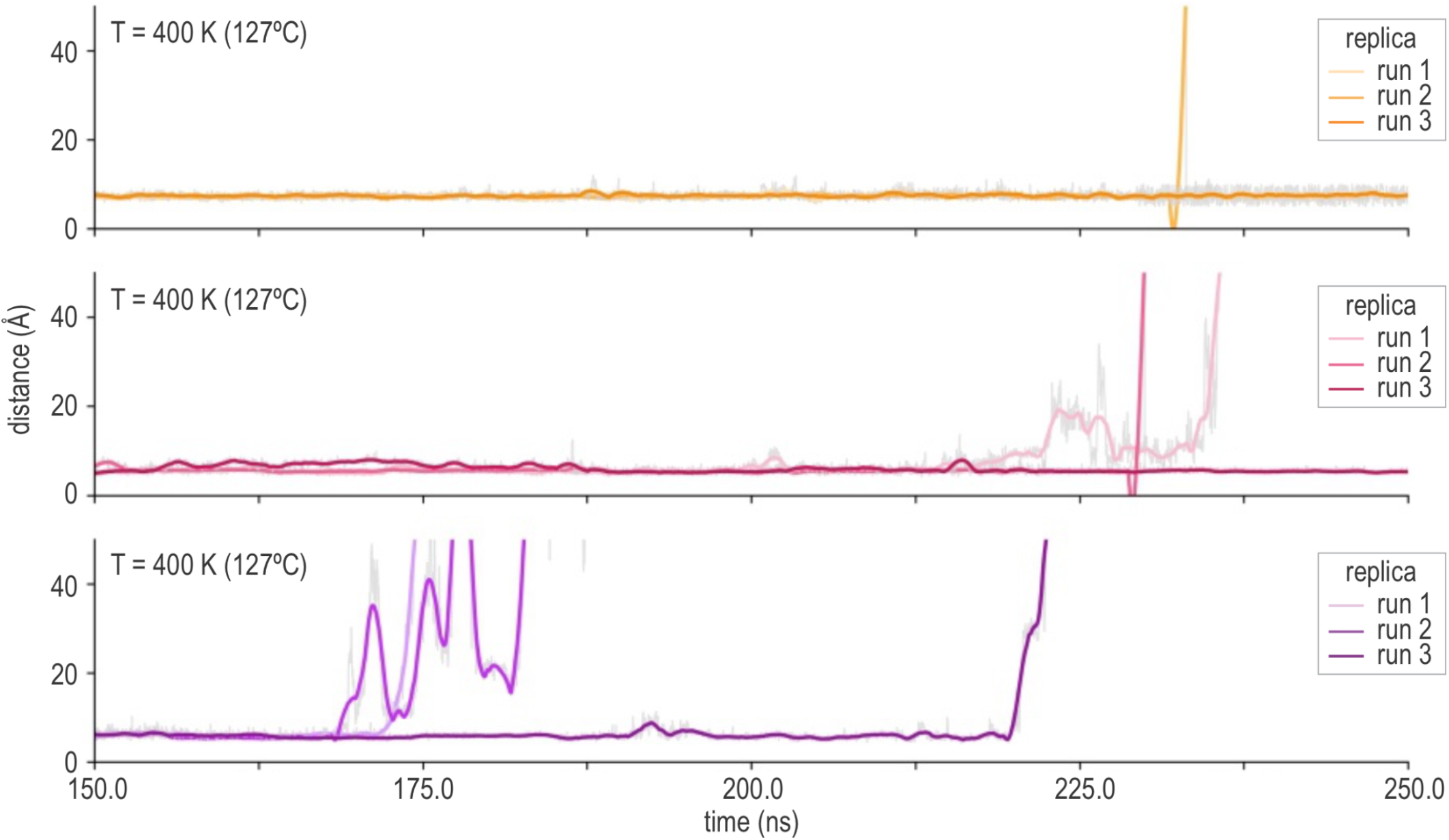
Accelerated thermal-stability test of NR4A3 binding site 1. Ligand-retention trajectories during the final 400 K phase of the accelerated stability simulations for PGA2 (orange, top), 20*α*-dihydrocortisol (pink, middle) and cortisol (purple, bottom) at binding site 1. Each panel shows the same ligand-to-site distance metric for three independent replicas from 150 to 250 ns; faint traces show instantaneous values and darker lines show the smoothed trajectories. These stress-test simulations extend the 300 K site-1 analysis summarized in Fig. 3k–o and discriminate the ligand-retention behaviour of the three complexes under a common accelerated perturbation. The 400 K phase is used as a comparative stability test rather than a physiological condition.

**Figure ED12:**
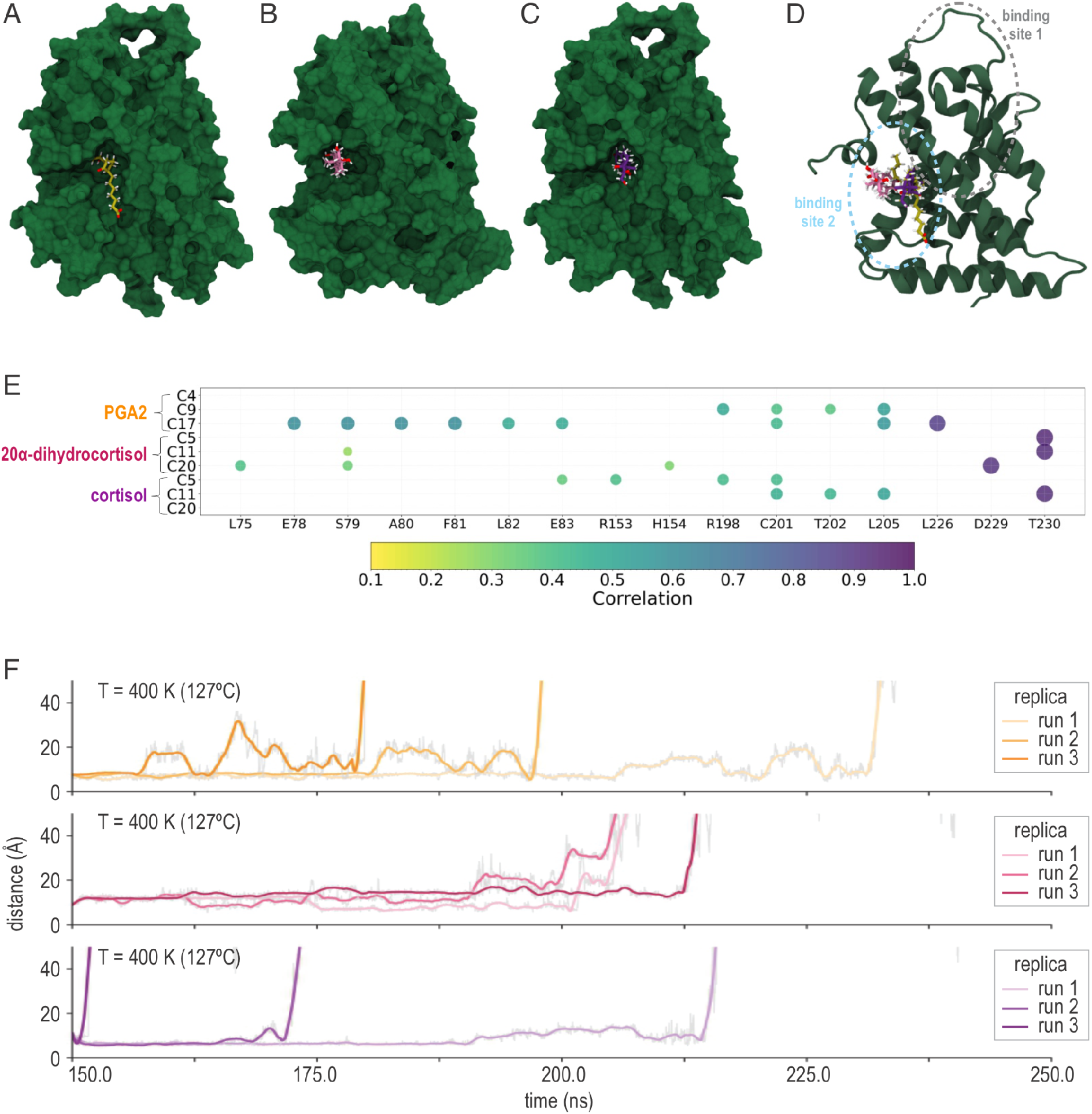
Complete analysis of the alternative NR4A3 binding site 2. **a–c**, NR4A3 molecular surface with PGA2 (**a**, orange), 20*α*-dihydrocortisol (**b**, pink) or cortisol (**c**, purple) positioned at the second candidate binding pocket. **d**, NR4A3 cartoon locating binding site 2 (blue dashed outline) relative to binding site 1 (grey dashed outline). **e**, Generalized correlations between site-2 residues and selected ligand atoms from the triplicate simulations; circle colour and size encode correlation strength from 0.1 (yellow) to 1.0 (purple). **f**, Accelerated 400 K ligand-retention trajectories for PGA2 (orange, top), 20*α*-dihydrocortisol (pink, middle) and cortisol (purple, bottom), with three independent replicas per ligand. Together, panels **a–f** provide the complete comparison with the more stable site-1 model used in Fig. 3k–o and Extended Data Fig. 11.

**Figure ED13:**
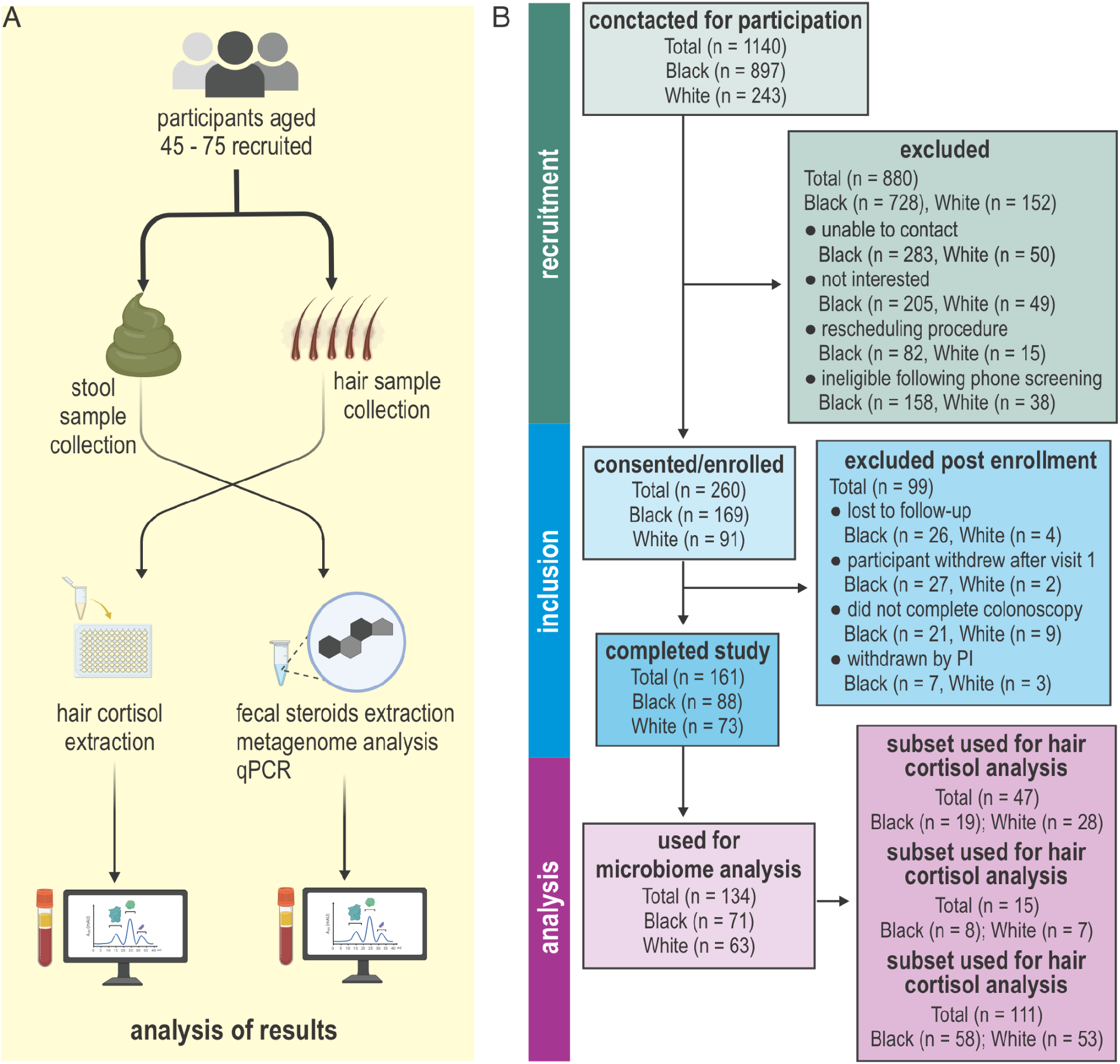
Human cohort sampling and analysis flow. **a**, Study overview. Adults aged 45–75 years undergoing screening or surveillance colonoscopy provided stool and hair samples; hair was used for cortisol quantification, whereas stool supported metagenomic, *desC* qPCR and steroid analyses. **b**, Recruitment and inclusion flow. Of 1,140 people contacted (897 Black and 243 White), 260 consented/enrolled and 161 completed the study (88 Black and 73 White). Analysis-specific subsets comprised 134 participants with faecal metagenomic data (71 Black, 63 White), 111 with faecal *desC* qPCR data (58 Black, 53 White), 47 with hair-cortisol measurements (19 Black, 28 White), and 15 participants with detectable faecal 20*α*-dihydrocortisol used for concentration comparisons (8 Black, 7 White). Reasons for exclusion before and after enrolment are enumerated in the flow chart; cohort characteristics for each assay subset are given in Supplementary Tables 3–6.

### Supplementary Information

### AKR1C1 vector sequence

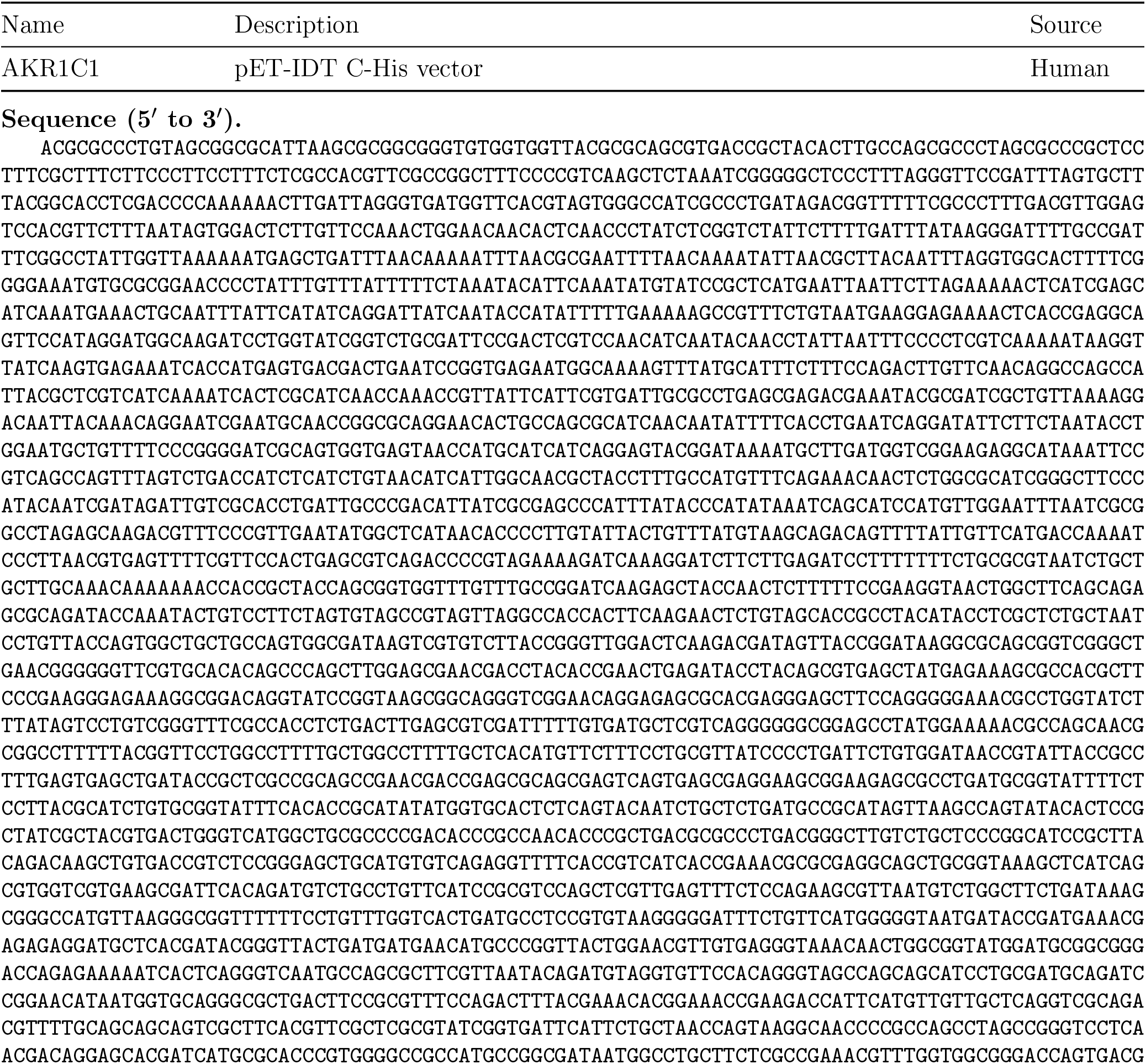
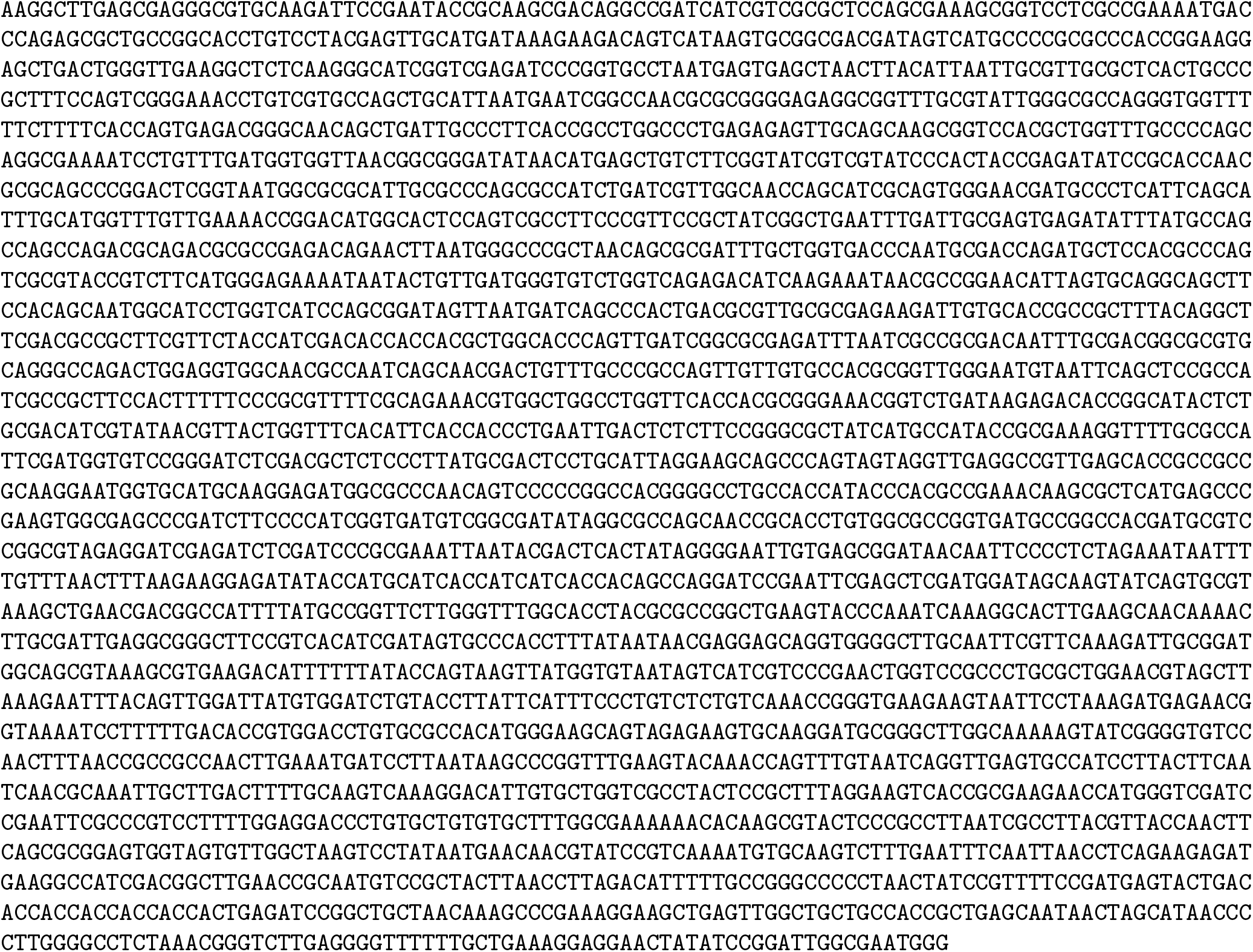

| Name | Description | Source |
| --- | --- | --- |
| AKR1C1 | pET-IDT C-His vector | Human |

## Supplementary tables

**Table S1:** Data collection and refinement statistics. Values in parentheses are for the highest-resolution shell unless otherwise indicated.

| Parameter | Value |
| --- | --- |
| PDB code | 4OH1 |
| Space group | I222 |
| Unit cell (Å) | $a = 63.87$, $b = 85.60$, $c = 149.09$ |
| Resolution (Å) | 29.4–2.0 (2.05–2.00) |
| Total reflections | 27,918 (1,995) |
| Completeness (%) | 99.5 (98.1) |
| Mean $I/\sigma(I)$ | 10.15 (1.4) |
| $R_{\text{merge}}$ | 0.05 (0.61) |
| $R_{\text{crystal}}/R_{\text{free}}$ | 0.168/0.216 (0.311, 0.297) |
| $R_{\text{free}}$ test set | 1,403 reflections (5.00%) |
| Wilson B-factor (Å<sup>2</sup>) | 35.5 |
| $F_o - F_c$ correlation (free) | 0.97 (0.95) |
| Total number of atoms | 2,918 |
| Average B, all atoms (Å<sup>2</sup>) | 49 |
| Ramachandran outliers | 0 |
| RMS bonds (Å) | 0.02 |
| RMS angles (°) | 1.8 |
| Sidechain outliers | 1.40% |
| RSRZ outliers | 7.90% |
| *Numbers in final model* |  |
| Residues | 351 |
| Protein atoms | 2,704 |
| Catalytic Zn<sup>2+</sup> | 2 |
| Water molecules | 193 |
| Chloride | 3 |
| 1,2-ethanediol | 1 |
| Tetraethylene glycol | 1 |
| *Mean B-values (Å<sup>2</sup>)* |  |
| Protein atoms | 36.2 |
| Tetraethylene glycol | 67.75 |
| 1,2-ethanediol | 72.75 |
| Zinc 1; zinc 2 | 47; 49 |
| Chloride 1; chloride 2 | 45; 57 |

**Table S2:** Elution profile of zinc-finger cysteine mutants by gel-filtration chromatography. The source PDF uses a second “Table S2” label; it is identified here as S2b for clarity.

| DesC mutant | $V_e/V_o$ | Estimated MW (kDa) |
| --- | --- | --- |
| C105A | 0.997871 | > 2000 |
| C108A | 0.997445 | > 2000 |
| C111A | 0.997658 | > 2000 |
| C119A | 0.998935 | > 2000 |
| C105A–C119A | 0.999148 | > 2000 |
| C108A–C119A | 0.999219 | > 2000 |
| WT | 1.0264 | 164 |

**Table S3:** Kinetic parameters for wild-type DesC and active-site mutants.

| Enzyme | $K_m$ ($\mu\text{M}$) | $V_{\text{max}}$ ($\text{nmol min}^{-1} \text{mg}^{-1}$) | $k_{\text{cat}}$ ($\text{min}^{-1}$) | $k_{\text{cat}}/K_m$ | Activity loss (%) |
| --- | --- | --- | --- | --- | --- |
| WT | $11.62 \pm 1.02$ | $2391.5 \pm 194.09$ | $92.31 \pm 7.49$ | $8.01 \pm 0.95$ | 0 |
| S47A | NA | NA | NA | NA | 100 |
| H50A | $23.99 \pm 0.26$ | $316.36 \pm 67.55$ | $12.5 \pm 0.89$ | $0.52 \pm 0.04$ | $71.81 \pm 6.02$ |
| S47T | $77.78 \pm 1.78$ | $16.66 \pm 0.90$ | $0.66 \pm 0.04$ | $0.0085 \pm 0.0006$ | $98.52 \pm 0.08$ |
| Y101R | $628.65 \pm 29.96$ | $44.80 \pm 7.54$ | $1.77 \pm 0.30$ | $0.0028 \pm 0.0006$ | $96.01 \pm 0.67$ |
| Y124R | $61.01 \pm 2.71$ | $204 \pm 3.45$ | $8.06 \pm 0.14$ | $0.13 \pm 0.01$ | $81.82 \pm 0.31$ |

**Table S4:** Secondary structural features of DesC wild type and mutants determined by circular dichroism spectroscopy.

| Protein | Helix | Strand | Turn | Unordered |
| --- | --- | --- | --- | --- |
| WT | 0.13 | 0.33 | 0.22 | 0.31 |
| S47A | $0.13 \pm 0.005$ | $0.32 \pm 0.008$ | $0.23 \pm 0.005$ | $0.32 \pm 0.006$ |
| H50A | $0.15 \pm 0.005$ | $0.30 \pm 0.005$ | $0.24 \pm 0.008$ | $0.31 \pm 0.006$ |
| S47T | $0.13 \pm 0.005$ | $0.33 \pm 0.006$ | $0.22 \pm 0.005$ | $0.32 \pm 0.006$ |
| Y101R | $0.14 \pm 0.01$ | $0.32 \pm 0.008$ | $0.23 \pm 0.006$ | $0.32 \pm 0.005$ |
| Y124R | $0.15 \pm 0.005$ | $0.31 \pm 0.005$ | $0.23 \pm 0.006$ | 0.30 |

**Table S5:** Demographics and lifestyle characteristics of the sequencing cohort by race.

| Characteristic | Black (n=69) | White (n=65) |
| --- | --- | --- |
| Age at recruitment, y (IQR) | 60 (55–63) | 60 (56–64) |
| Sex, n M/F | 26/43 | 36/29 |
| *Highest level of education, n (%)* |  |  |
| Some high school | 3 (4) | 0 |
| High school graduate/GED | 12 (17) | 2 (3) |
| Some college/Associate degree | 37 (54) | 11 (17) |
| Completed 4-year college | 7 (10) | 22 (34) |
| Graduate school | 10 (14) | 30 (46) |
| *Annual household income, n (%)* |  |  |
| < \$10,000 | 8 (12) | 1 (2) |
| \$10,000–\$19,999 | 7 (10) | 2 (3) |
| \$20,000–\$49,999 | 18 (26) | 6 (9) |
| \$50,000–\$100,000 | 17 (25) | 9 (14) |
| \$100,000+ | 18 (26) | 46 (72) |
| Smoking status, n Y/N | 14/55 | 4/61 |
| BMI, kg m<sup>-2</sup> (IQR) | 33.6 (28.05–38.2) | 28.90 (24.45–32.05) |
| NSAID use, n Y/N | 34/34 | 41/23 |

**Table S6:**
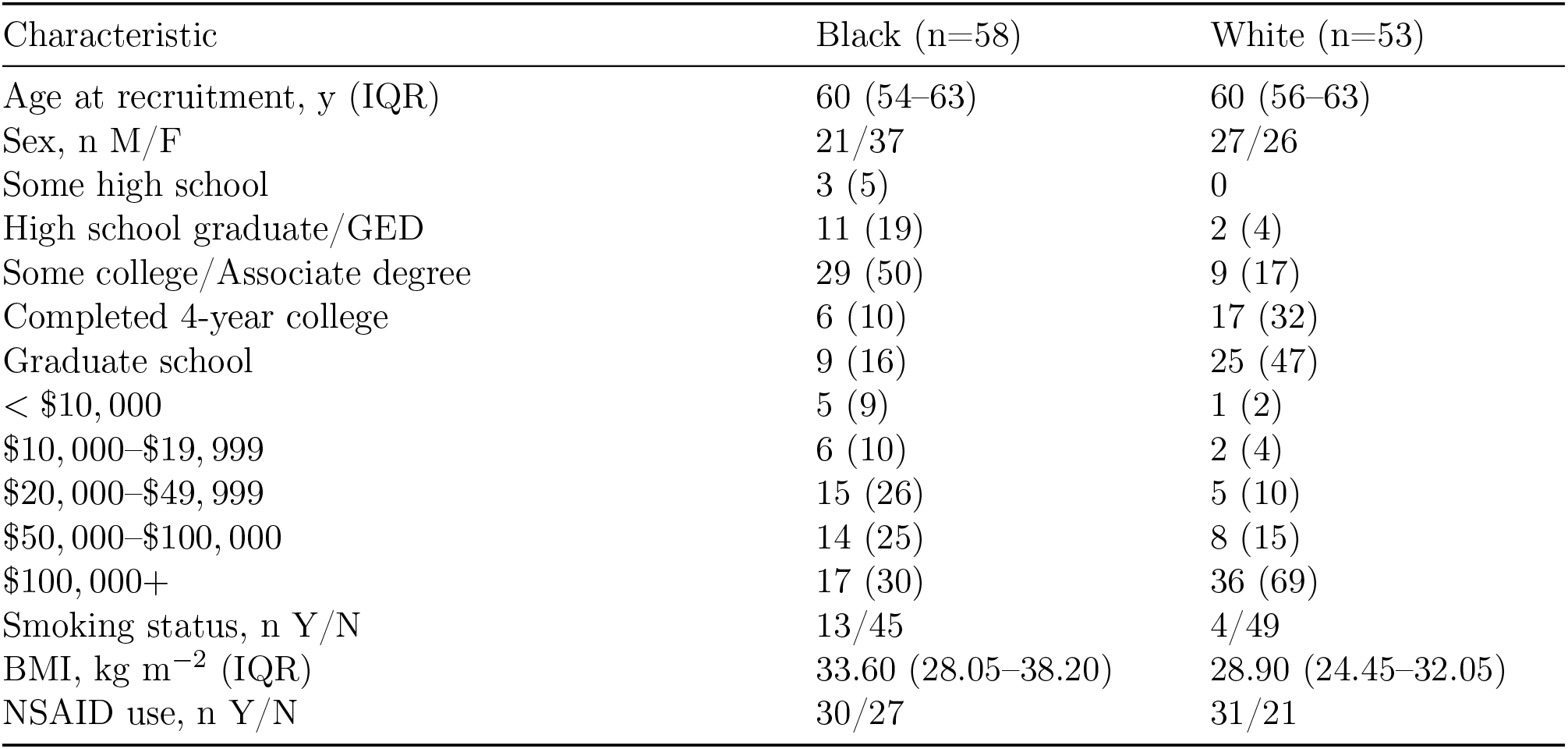
Demographics and lifestyle characteristics of the qPCR cohort by race.

| Characteristic | Black (n=58) | White (n=53) |
| --- | --- | --- |
| Age at recruitment, y (IQR) | 60 (54–63) | 60 (56–63) |
| Sex, n M/F | 21/37 | 27/26 |
| Some high school | 3 (5) | 0 |
| High school graduate/GED | 11 (19) | 2 (4) |
| Some college/Associate degree | 29 (50) | 9 (17) |
| Completed 4-year college | 6 (10) | 17 (32) |
| Graduate school | 9 (16) | 25 (47) |
| < \$10,000 | 5 (9) | 1 (2) |
| \$10,000–\$19,999 | 6 (10) | 2 (4) |
| \$20,000–\$49,999 | 15 (26) | 5 (10) |
| \$50,000–\$100,000 | 14 (25) | 8 (15) |
| \$100,000+ | 17 (30) | 36 (69) |
| Smoking status, n Y/N | 13/45 | 4/49 |
| BMI, kg m<sup>-2</sup> (IQR) | 33.60 (28.05–38.20) | 28.90 (24.45–32.05) |
| NSAID use, n Y/N | 30/27 | 31/21 |

**Table S7:** Demographics and lifestyle characteristics of the cohort with hair cortisol values by race.

| Characteristic | Black (n=19) | White (n=28) |
| --- | --- | --- |
| Age at recruitment, y (IQR) | 61 (57–63) | 61 (54–66) |
| Sex, n M/F | 4/15 | 12/16 |
| Some high school | 2 (10.5) | 0 |
| High school graduate/GED | 2 (10.5) | 1 (4) |
| Some college/Associate degree | 11 (58) | 6 (21) |
| Completed 4-year college | 1 (5) | 10 (36) |
| Graduate school | 3 (16) | 11 (39) |
| < \$10,000 | 3 (16) | 0 |
| \$10,000–\$19,999 | 2 (11) | 1 (4) |
| \$20,000–\$49,999 | 5 (26) | 2 (7) |
| \$50,000–\$100,000 | 4 (21) | 4 (14) |
| \$100,000+ | 5 (26) | 21 (75) |
| Smoking status, n Y/N | 5/14 | 1/27 |
| BMI, kg m<sup>-2</sup> (IQR) | 35.20 (31.30–41.20) | 28.25 (23.55–30.75) |
| NSAID use, n Y/N | 11/8 | 22/6 |
| Hair cortisol, pg/mg (IQR) | 8.6 (4.8–34.2) | 2.4 (1.4–4.1) |

**Table S8:** Demographics and lifestyle characteristics of participants with detectable fecal steroid by race.

| Characteristic | Black (n=8) | White (n=7) |
| --- | --- | --- |
| Age at recruitment, y (IQR) | 61 (59–68) | 56 (51–69) |
| Sex, n M/F | 1/7 | 3/4 |
| Some high school | 0 | 0 |
| High school graduate/GED | 1 (12.5) | 0 |
| Some college/Associate degree | 4 (50) | 2 (28.5) |
| Completed 4-year college | 1 (12.5) | 3 (43) |
| Graduate school | 2 (25) | 2 (28.5) |
| < \$10,000 | 1 (12.5) | 0 |
| \$10,000–\$19,999 | 1 (12.5) | 0 |
| \$20,000–\$49,999 | 3 (37.5) | 1 (14) |
| \$50,000–\$100,000 | 1 (12.5) | 0 |
| \$100,000+ | 2 (25) | 6 (86) |
| Smoking status, n Y/N | 3/5 | 0/7 |
| BMI, kg m<sup>-2</sup> (IQR) | 34.65 (31.93–35.8) | 25.10 (22.00–27.90) |
| NSAID use, n Y/N | 4/4 | 6/1 |

**Table S9:**
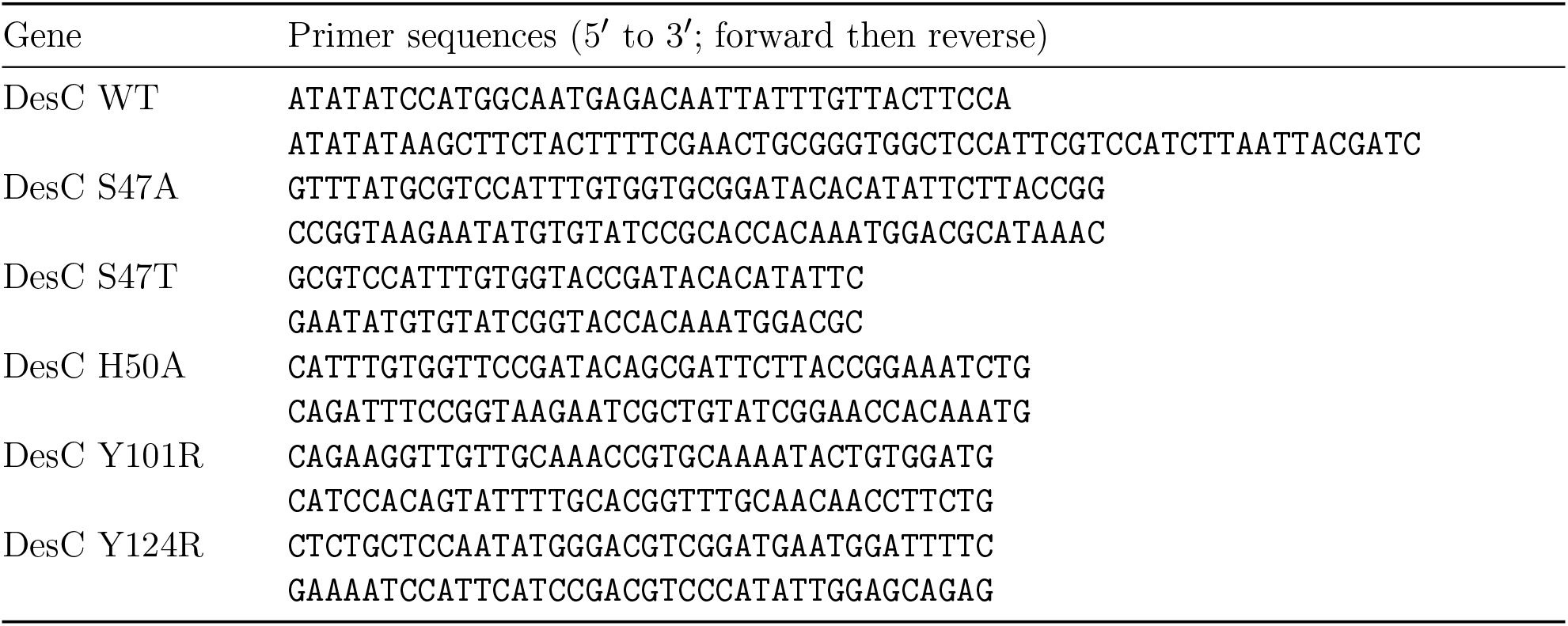
Primers employed in the study.

**Table S10:** Plasmids used in this study.

| Name | Description | Reference |
| --- | --- | --- |
| pCas9 | Temperature-sensitive helper plasmid expressing *S. pyogenes* Cas9 and λ-Red recombinase; kanamycin resistance | Addgene #62225 |
| pTargetF | pMB1-based gRNA expression plasmid for locus-specific targeting; streptomycin resistance | Addgene #62226 |
| pTarget(SS3) | pTargetF carrying a gRNA cassette targeting the SS3 intergenic locus in *E. coli* BL21(DE3) | This study |
| pTarget(SS3)_desCWT | pTarget(SS3) carrying the wild-type *desC* expression cassette and SS3 homology arms | This study |
| pTarget(SS3)_desCM | pTarget(SS3) carrying the S47A mutant *desC* expression cassette and SS3 homology arms | This study |

**Table S11:**
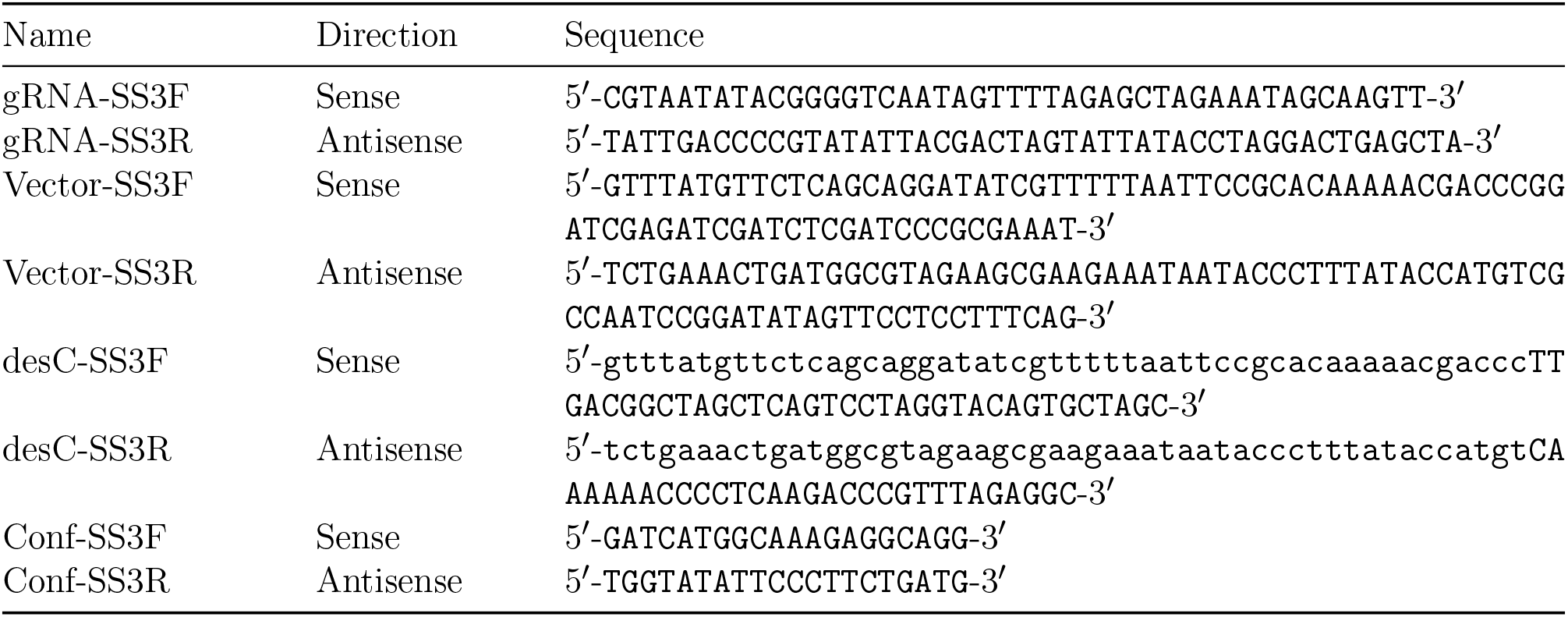
Primers used for CRISPR engineering.

**Table S10.**
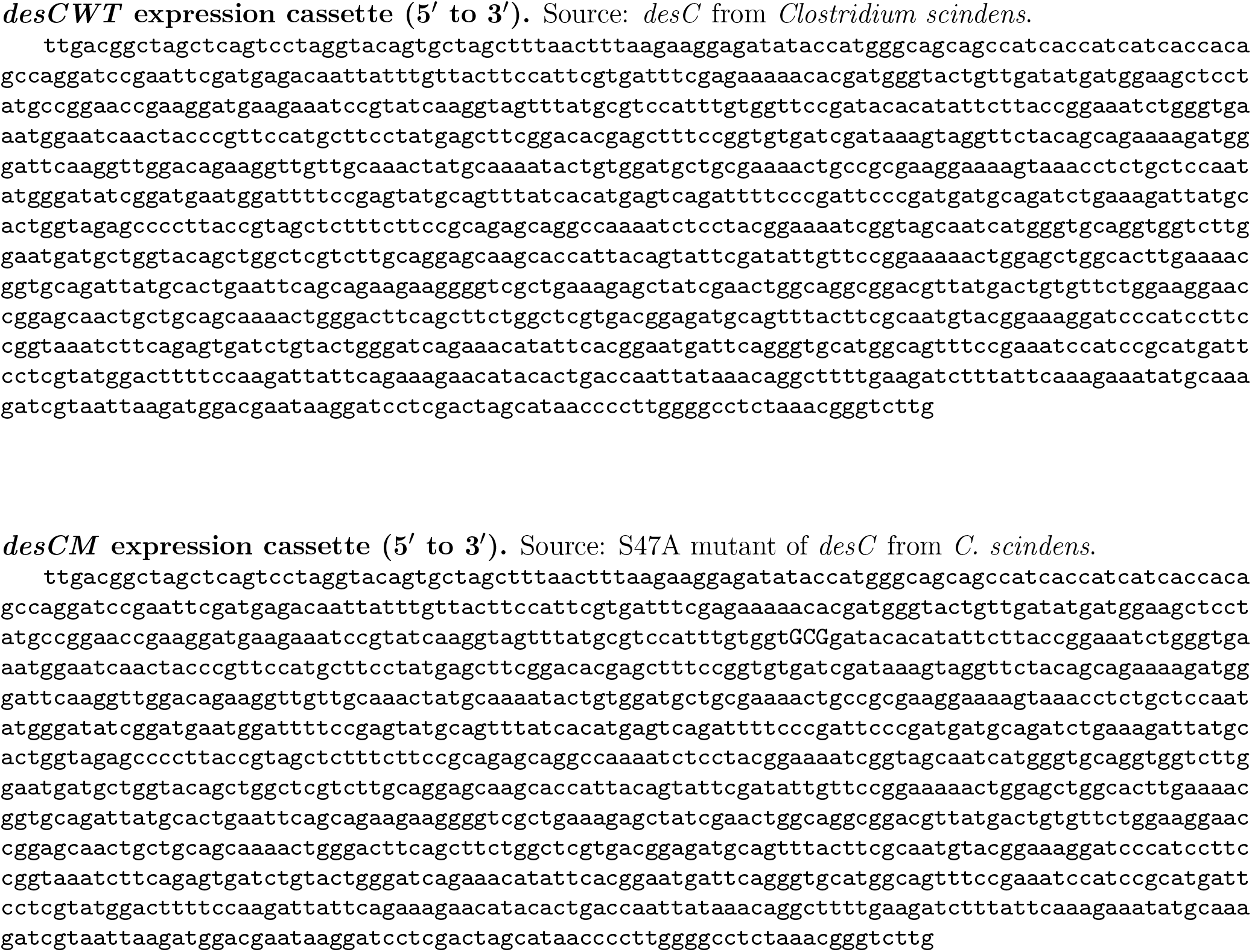
Synthetic DNA sequences used in this study.

**Table S12:**
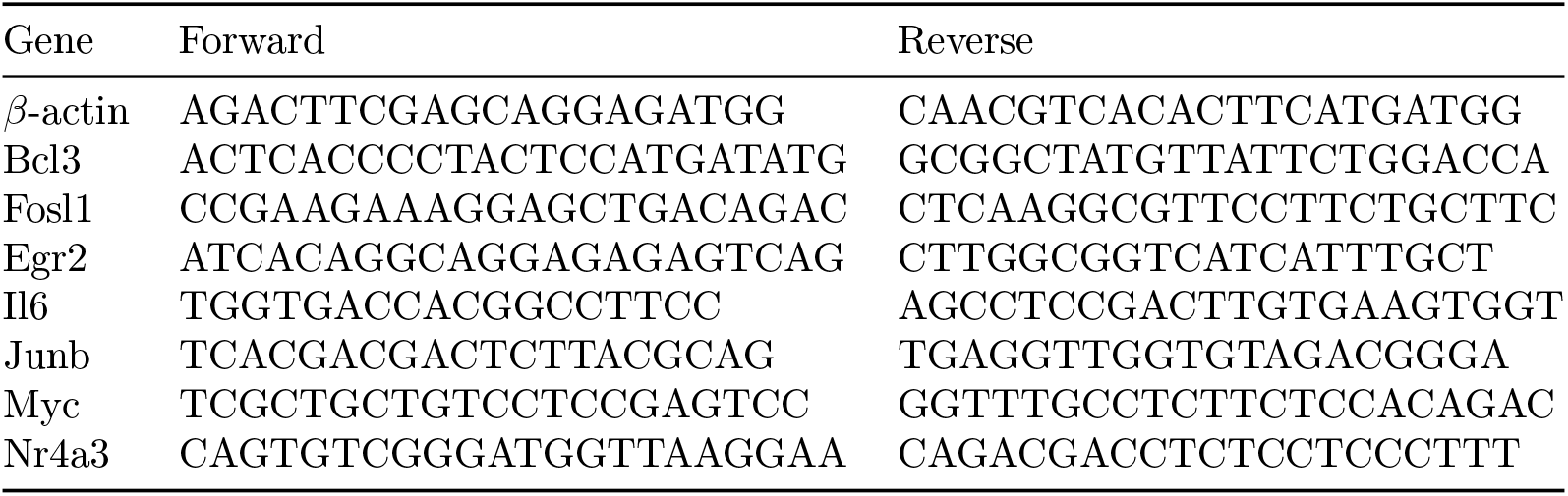
Primers used for qPCR detection in intestinal epithelial cells after *Nr4a3* knockdown with siRNA.

**Table S13:**
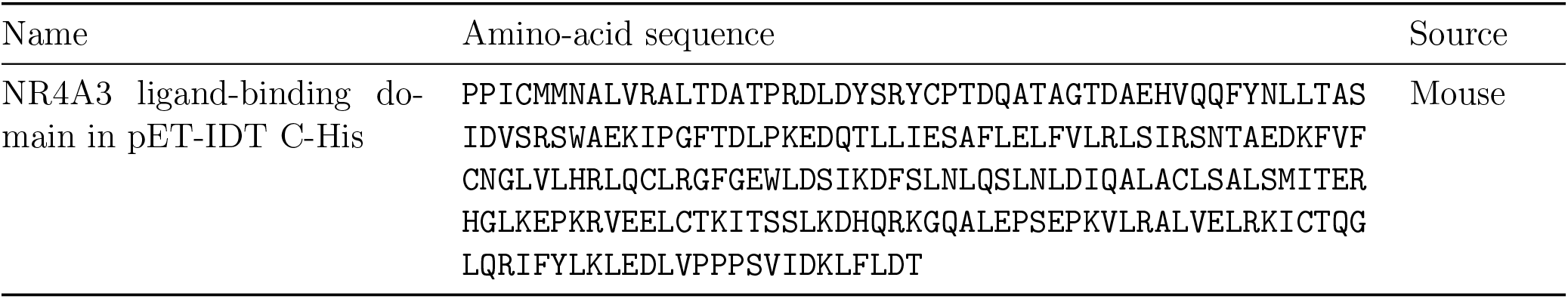
Mouse NR4A3 ligand-binding-domain amino-acid sequence.

**Table S14:**
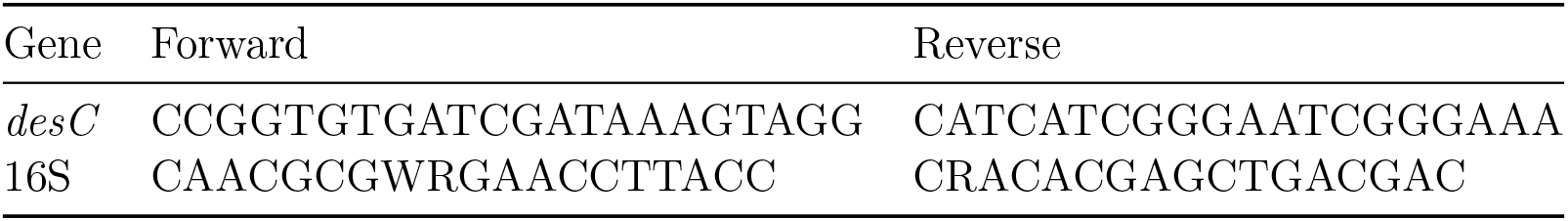
Primers used for qPCR detection in human fecal samples.

## Supplementary materials and methods

### Bacterial strains, steroids and reagents

*Clostridium scindens* ATCC 35704 was maintained as glycerol stocks at *−*80 *^◦^*C. *Escherichia coli* DH5*α* was used for routine plasmid propagation, BL21 CodonPlus (DE3)-RIPL for recombinant DesC production and BL21(DE3) for construction of the strains used in the gnotobiotic experiment. The pET-51b and pETDuet-1 vectors were obtained commercially. Restriction enzymes and Gibson Assembly reagents were from New England Biolabs. Strep-Tactin resin was from IBA Lifesciences. Cortisol and 20*α*-dihydrocortisol standards were obtained from Steraloids. Unless otherwise indicated, reagents were analytical grade.

### DesC cloning, expression and purification

The open reading frame encoding DesC (EDS07887) was amplified from *C. scindens* genomic DNA with the primers in Supplementary Table 7 and cloned into pET-51b as described previously [1]. Insert identity was confirmed by Sanger sequencing. Recombinant plasmid was transformed into *E. coli* BL21 CodonPlus (DE3)-RIPL. Colonies were grown in LB with ampicillin (100 *µ*g ml*^−^*^1^) and chloramphenicol (50 *µ*g ml*^−^*^1^); 1-litre cultures were induced at OD_600_ *≈* 0.3 with 0.1 mM IPTG and then grown for 16 h at 16 °C. Cells were collected at 4,000*×g* for 30 min at 4 °C, resuspended in 20 mM Tris-HCl, 150 mM NaCl, 20% glycerol and 10 mM 2-mercaptoethanol, pH 7.9, and lysed by four passes through an EmulsiFlex C-3 homogenizer. Clarified lysate was obtained at 20,000*×g* for 30 min at 4 °C. DesC was purified on Strep-Tactin resin and eluted in the same buffer containing 2.5 mM d-desthiobiotin. Purity was assessed by SDS–PAGE and Coomassie staining.

### DesC crystallization and structure determination

Selenomethionine-labelled DesC was screened against the JCSG Core Suite. Apo-DesC crystals were obtained by sitting-drop vapour diffusion at 20 °C against 0.2 M MgCl_2_, 24% PEG 400 and 0.1 M HEPES, pH 6.75. Crystals were supplemented with 10% 1,2-ethanediol, harvested and flash-frozen in liquid nitrogen. Diffraction data were collected at beamline BL14-1 of the Stanford Synchrotron Radiation Light-source. Data were indexed and integrated with XDS and scaled with XSCALE [2, 3]. Selenium multiwavelength anomalous-dispersion phasing was performed with SHELX and the model was refined with Refmac [4–7]. The final coordinates and structure factors are deposited as PDB 4OH1. Data-collection and refinement statistics are given in Supplementary Table 2a, with representative zinc-site electron density shown in Extended Data Fig. 2.

### DesC site-directed mutagenesis and structural controls

Candidate catalytic, substrate-recognition and structural-zinc residues were selected from the DesC structure, sequence alignments and the MD-derived substrate pose. Variants were produced in the pET-51b DesC construct with the QuikChange Lightning kit and the primers listed in Supplementary Table 7. Plasmids were propagated in XL10 cells, sequence-verified, and mutant proteins were expressed and purified with the same protocol used for wild-type DesC.

Circular-dichroism spectra were collected on a J-815 spectropolarimeter using 0.2 mg ml*^−^*^1^ protein in 10 mM KH_2_PO_4_, pH 7.5. Spectra were recorded at 25 °C from 190 to 260 nm in a 0.1-cm quartz cuvette at 50 nm min*^−^*^1^, using a 0.1-nm pitch and five accumulations. Secondary-structure fractions were estimated with CDSSTR using reference set 4 on the DICHROWEB server. Values are summarized in Supplementary Table 2d and representative Y101R/Y124R spectra are shown in Extended Data Fig. 4. Structural-zinc variants were additionally examined by gel-filtration chromatography; elution profiles are reported in Supplementary Table 2b.

### DesC enzyme kinetics and substrate profiling

DesC activity was quantified by following NADH oxidation at 340 nm (*ε* = 6, 220 M*^−^*^1^cm*^−^*^1^). Standard reactions contained 50 mM phosphate buffer, pH 7.5, 50 *µ*M cortisol, 150 *µ*M NADH and 0.05 *µ*M enzyme in a final volume of 0.5 ml. Reactions were initiated by adding enzyme. Initial velocities measured across substrate concentrations were fitted to the Michaelis–Menten equation by nonlinear regression in GraphPad Prism. Kinetic parameters for wild-type DesC and active-site/substrate-pocket variants are given in Supplementary Table 2c.

Substrate-specificity measurements used 200 *µ*M steroid and three biological replicates per substrate. Activities were normalized to prednisone for DesC. Steroid assignments were verified against authentic standards by chromatography (Extended Data Fig. 1). Values reported in Fig. 1b represent mean*±*s.d.

### DesC isothermal titration calorimetry

ITC measurements were performed with a MicroCal VP-ITC instrument with a 1.4-ml sample cell. Proteins were dialysed against 50 mM sodium phosphate, 150 mM NaCl and 10% glycerol, pH 7.5, and NADH and cortisol were prepared in matched buffer. DesC proteins (50 *µ*M), with or without pyridine nucleotide, were titrated with 28 successive 10-*µ*l aliquots of 0.5 mM cortisol at 300-s intervals. Data were fitted to a single-site nonlinear-regression model with MicroCal Origin. Thermodynamic quantities were related by Δ*G* = Δ*H − T* Δ*S* = *−RT* ln *K_a_*. Experiments used to evaluate Tyr101/Tyr124-mediated recognition are shown in Extended Data Fig. 4.

### Human AKR1C1 production and activity

Human *AKR1C1*, encoding 20*α*-hydroxysteroid dehydrogenase, was codon-optimized for *E. coli* and expressed with an N-terminal His tag from a pET-IDT vector (Supplementary Table 1 and Extended Data Fig. 1). Protein was produced in BL21 cells after IPTG induction, cells were disrupted with an Emulsi-Flex C-3 homogenizer, and AKR1C1 was purified on a 5-ml HisTrap column using an AKTAxpress FPLC system. Activity was measured by NADPH oxidation at 340 nm in 100 mM potassium phosphate, pH 7.0, 200 *µ*M steroid, 200 *µ*M NADPH and 1 *µ*M enzyme in 0.2 ml. Absorbance was measured every 8 s for 8 min. Initial velocities were normalized to progesterone, defined as 100% activity.

### Construction of the DesC–NADH–cortisol model

BLAST searches identified cofactor-bound structural homologues with PDB codes 4ILK, 4EJ6, 4A2C, 3QE3, 3GFB, 2DQ4, 2DFV, 2D8A, 1PL7 and 1E3J. Structures were aligned to apo-DesC in VMD [8], and their bound cofactors were used to define the starting NADH geometry (Extended Data Fig. 3). Cortisol was positioned with its C20 carbonyl oriented towards the NADH hydride donor and catalytic zinc, and the local ligand–protein geometry was relaxed with QwikMD [9]. Five independent model-building and refinement replicas were generated. All five converged to a similar, stable arrangement of NADH and cortisol within the DesC active site, providing a reproducible starting geometry for subsequent simulations. These models were then subjected to independent classical molecular dynamics simulations.

### Classical molecular dynamics of DesC

The DesC–NADH–cortisol complexes were prepared with VMD and QwikMD, solvated in explicit TIP3P water and simulated with NAMD under periodic boundary conditions [8–10]. CHARMM36 parameters were used for the biomolecular environment. Simulations were performed in the NpT ensemble at 300 K and 1 bar with Langevin temperature and pressure coupling. Short-range nonbonded interactions used a 12-Å cutoff and long-range electrostatics were treated with particle-mesh Ewald [11]. The r-RESPA multiple-time-step scheme updated van der Waals interactions every step and long-range electrostatics every second step [12].

Five independent MD replicas were performed. Each replica was subjected to 5,000 minimization steps followed by 10 ns of equilibration with restraints on the protein backbone and ligand non-hydrogen atoms and a subsequent 100-ns unrestrained equilibrium simulation. Across the five replicas, the DesC–NADH–cortisol complex converged to a consistent active-site configuration, with comparable positioning of the cortisol C20 carbonyl relative to NADH and the catalytic centre. Analysis of the independent trajectories further identified Tyr101 and Tyr124 as persistent cortisol-recognition residues. A representative equilibrated configuration from this reproducible conformational ensemble was selected to initiate the QM/MM calculations.

### Hybrid QM/MM simulations of DesC catalysis

Electrostatically embedded QM/MM calculations used the NAMD QM/MM interface coupled to MOPAC [13, 14]. The MM region used CHARMM36. The 573-atom quantum region used the semiempirical PM7 Hamiltonian and contained cortisol, NADH, the catalytic zinc and the surrounding active-site environment required to represent the reaction electronically. Thirty-six QM–MM covalent boundaries were treated with link atoms using the charge-shift approach, with the corresponding redistribution of nearby classical charges [13, 15].

The QM/MM system was initiated from a representative configuration of the conformational state consistently sampled across the five independent classical MD replicas. A 20-ps unbiased QM/MM equilibration was first performed using a 2-fs integration step. Candidate reaction mechanisms were subsequently probed with QM/MM steered MD using a 0.5-fs step. Bias was applied only to the initiating bond rearrangements; subsequent proton transfers were allowed to occur without direct forcing when energetically favourable. These calculations identified a mechanism without an intervening catalytic water, in which hydride is transferred from NADH to the carbonyl carbon of cortisol C20 while Ser47 protonates the carbonyl oxygen. Proton transfer from the NAD^+^ ribose to Ser47 and from His50 to the ribose completes the relay. The resulting reaction trajectory provided the initial pathway for subsequent finite-temperature string optimization and free-energy sampling.

### String optimization and free-energy sampling

The QM/MM steered-MD trajectory supplied the initial reaction path for finite-temperature string optimization [16]. Collective variables were interatomic donor–hydrogen and hydrogen–acceptor distances describing hydride transfer and the subsequent three proton transfers; their progression is shown in Extended Data Fig. 5. The path was represented by 25 images. Independent short QM/MM trajectories were launched around each image to sample the local configurational ensemble, and the resulting mean positions were iteratively used to update and reparameterize the string until the reaction coordinate converged. In aggregate, 750 QM/MM trajectories were propagated per string iteration, giving 4.5 ns of QM/MM sampling for pathway optimization.

Free energy along the optimized path was estimated with extended adaptive biasing force (eABF), using the path variables *S*, describing progression along the string, and *Z*, describing displacement perpendicular to it [17, 18]. Multiple walkers were distributed along the reaction path, with 10,000 MD steps per walker. Production sampling accumulated approximately 3 ns of additional QM/MM dynamics across the distributed calculation. Thus, although the string optimization and eABF calculation were performed for a single reaction pathway, the resulting mechanism and free-energy profile derive from ensemble-based enhanced sampling comprising many independently propagated QM/MM trajectories rather than from a single transition trajectory. The resulting profile reports the chemical transformation along the selected coordinate and therefore does not include substrate association, large-scale conformational rearrangements preceding chemistry or product release.

### NR4A3 model, ligand preparation and docking

Full-length mouse NR4A3 was modelled with AlphaFold 3. The selected model had a combined ipTM+pTM score of 0.88. PGA2, 20*α*-dihydrocortisol and cortisol were obtained from PubChem and protonated for pH 7.4 with OpenBabel. Ligand binding was evaluated at two candidate sites, with three independent replicas generated for each ligand–site combination. Thus, the complete design comprised six ligand–site combinations, each evaluated in triplicate.

At site 1, docking scores were *−*5.42, *−*6.42 and *−*6.37 kcal mol*^−^*^1^ for PGA2, 20*α*-dihydrocortisol and cortisol, respectively. At site 2, scores were *−*6.2 and *−*6.1 kcal mol*^−^*^1^ for 20*α*-dihydrocortisol and cortisol; PGA2 was positioned using its experimentally observed binding geometry from PDB 5Y41 rather than redocked. Site 1 corresponds to the structures and poses shown in Fig. 3k–n, whereas the complete site-2 comparison is shown in Extended Data Fig. 11.

### NR4A3 molecular dynamics and correlation analysis

Each ligand–NR4A3 complex was prepared with VMD/QwikMD, solvated in an explicit TIP3P water box, neutralized and adjusted to 0.15 M NaCl. Simulations were performed with NAMD 3 under periodic boundary conditions. Systems underwent 1,000 minimization steps followed by gradual heating from 10 to 300 K in 1-K increments and 0.5 ns of NpT equilibration at 300 K and 1 atm. Production comprised an initial 5-ns NpT phase with stabilizing extra bonds, a 50-ns unrestrained NpT phase at 300 K, a 100-ns annealing phase in which the temperature increased linearly from 300 to 400 K at 1 K ns*^−^*^1^, and a final 100-ns phase at 400 K.

All six ligand–site combinations—three ligands at each of the two candidate binding sites—were simulated in three independent replicas, yielding 18 independent MD trajectories in total. Each replica underwent the complete 50-ns unrestrained production, 100-ns annealing and 100-ns 400-K phases. The high-temperature portion was used as a comparative ligand-retention stress test rather than to estimate binding affinity. Site-1 400-K trajectories are shown in Extended Data Fig. 10, whereas the corresponding site-2 trajectories are shown in Extended Data Fig. 11.

Ligand retention was quantified using consistent site-specific distance metrics across the three independent replicas. Generalized correlations between selected ligand atoms and receptor residues were calculated across the trajectories to identify reproducible ligand-specific patterns of dynamic communication. The site-1 map shown in Fig. 3o and the site-2 map in Extended Data Fig. 11 were generated using the same analysis framework and incorporated the replicate trajectories. Docking and MD were used to identify, test and compare candidate binding modes, whereas ligand affinity was determined independently by ITC.

### NR4A3 ligand-binding-domain production and ITC

The mouse NR4A3 ligand-binding-domain construct is shown in Extended Data Fig. 12 and its amino-acid sequence in Supplementary Table 14. The codon-optimized sequence was expressed from a pET-IDT C-His vector in BL21 *E. coli* after IPTG induction. Cells were collected for 40 min at 4,000 r.p.m. and 4 °C, disrupted with an EmulsiFlex C-3 homogenizer and purified on a 5-ml HisTrap column with an AKTAxpress system. A band of approximately 26 kDa was verified by SDS–PAGE.

ITC was performed with a MicroCal PEAQ-ITC instrument with a 0.28-ml cell. Protein was dialysed against 50 mM sodium phosphate, 150 mM NaCl and methyl-*β*-cyclodextrin, pH 7.5, and PGA2, 20*α*-dihydrocortisol and cortisol were prepared in matched buffer. NR4A3 ligand-binding domain (100 *µ*M) was titrated with 13 successive 3-*µ*l aliquots of 2 mM ligand at 150-s intervals. Isotherms were fitted to a single-site model in MicroCal PEAQ-ITC software. Three independently prepared protein samples were analysed for each ligand.

### CRISPR–Cas9 plasmid construction

The pCas plasmid (Addgene #62225) supplied Cas9 and *λ*-Red recombinase, whereas pTargetF (Addgene #62226) supplied the guide-RNA cassette [19]. To target the SS3 intergenic locus between *ompW* and *yciE*, the pTargetF guide was replaced by an SS3-specific guide using the primers in Supplementary Table 9. Circular donor DNA was assembled in pTarget by Gibson assembly using 50-bp homology arms flanking SS3 [20]. The constitutive expression cassette contained the J23100 promoter, pET ribosome-binding site, His tag, the relevant *desC* allele and T7 terminator. Wild-type and S47A donor sequences are given in Supplementary Table 10. Constructs were verified by restriction digestion and sequencing.

Before *desC* integration, the SS3 locus and expression architecture were validated with EGFP (Extended Data Fig. 6). Constitutive J23100-pETRBS-EGFP and inducible T7-pETRBS-EGFP cassettes were compared episomally and after chromosomal insertion. For fluorescence measurements, overnight cultures were diluted to OD_600_ = 0.1, grown for 12 h at 25 °C and, where applicable, induced with 0.5 mM IPTG. Cells were washed in PBS, adjusted to equal OD_600_, transferred to black 96-well plates and measured at 485-nm excitation/512-nm emission.

### Construction and validation of Eco_*desC* WT and Eco_*desC* M

pCas was electroporated into *E. coli* BL21(DE3), and *λ*-Red recombination was induced with L-arabinose. pTarget(SS3) carrying either the wild-type *desC* donor or the S47A donor was then introduced. Transformants were selected on LB with kanamycin (50 *µ*g ml*^−^*^1^) and streptomycin (50 *µ*g ml*^−^*^1^); integration was screened by colony PCR with Conf-SS3F/Conf-SS3R and sequence verification. pTarget was cured after growth in kanamycin plus 0.5 mM IPTG and identified by streptomycin sensitivity. pCas was then removed by overnight growth at 37 °C without selection. The resulting strains are Eco_*desC* WT and Eco_*desC* M.

To verify function, strains were precultured overnight, diluted into 25 ml LB containing 50 *µ*M cortisol and 5 g l*^−^*^1^ glycerol at OD_600_ = 0.1, and grown at 37 °C. Cell density was measured spectrophotometrically. Glycerol was quantified on an Agilent 1200 HPLC with a refractive-index detector and Rezex ROA-Organic Acid H+ column using 0.005 N H_2_SO_4_ at 0.6 ml min*^−^*^1^ and 50 °C. Cortisol and 20*α*-dihydrocortisol were measured on a Shimadzu LC-20A HPLC with diode-array detection and a CAPCELL PAK C18 MG column at 30 °C; 50% methanol was used at 0.3 ml min*^−^*^1^, with detection at 254 nm.

### Gnotobiotic mouse experiment

Animal procedures were approved by the University of Illinois Urbana–Champaign IACUC (protocol 19222). Germ-free C57BL/6J mice were maintained in flexible-film isolators in the Rodent Gnotobiotic Facility and monitored by aerobic/anaerobic culture, Gram staining and qPCR [21]. Mice were transferred aseptically into sealed positive-pressure cages before experimentation. Engineered *E. coli* cultures were grown from single colonies for 16 h at 37 °C, washed in sterile PBS and prepared for oral gavage. Mice received 10^8^ colony-forming units of Eco_*desC* WT or Eco_*desC* M by oral gavage on days 1 and 3. From day 4 through day 7, the indicated groups received sterile drinking water containing cortisol at 10 mg kg*^−^*^1^ day*^−^*^1^; controls received sterile water. Mice were euthanized on day 7. Serum, caecal contents and intestinal epithelial scrapings were collected. Four mice per strain-by-treatment group are represented in the reported metabolomic and transcriptomic analyses in Fig. 2 and Extended Data Fig. 7.

### Mouse steroid extraction and LC–MS/MS

Caecal material was dispersed in 0.3 ml deionized water and supplemented with 0.1 ml internal-standard solution. Serum aliquots (0.1 ml) were mixed with 0.1 ml water and 0.1 ml internal-standard solution. Samples were loaded onto Biotage Isolute supported-liquid-extraction columns and allowed to absorb for 5 min. Steroids were eluted with 1.75 ml methyl-*tert*-butyl ether, dried under vacuum and reconstituted in 0.2 ml 40:60 methanol:water. Aliquots were separated by two-dimensional liquid chromatography. The first dimension used an Agilent 1260 HPLC with a Hypersil GOLD C4 cleanup column; analytes were then switched onto a Kinetex biphenyl C18 analytical column and resolved on an Agilent 1290 UPLC with water/methanol gradients containing 0.2 mmol l*^−^*^1^ ammonium fluoride. Eluate entered an Agilent 6495A triple-quadrupole mass spectrometer operating in positive electrospray-ionization mode with dynamic multiple-reaction monitoring.

### 16S rRNA gene sequencing of mouse samples

Genomic DNA was extracted from approximately 0.05 g caecal material with the QIAamp PowerFecal Pro DNA kit. The V4 region of the bacterial 16S rRNA gene was sequenced as 2*×*250-bp paired-end reads on an Illumina MiSeq at the Roy J. Carver Biotechnology Center. Reads were demultiplexed with bcl2fastq v2.20, quality-checked with FastQC and analysed in QIIME 2 v2023.7 [22]. Genus-level profiles are shown in Extended Data Fig. 7.

### Primary mouse intestinal epithelial cells

Primary intestinal epithelial cells were isolated from conventionally raised C57BL/6 mice after opening and washing the intestine, enzymatic digestion and recovery of epithelial material. Cells were maintained in complete DMEM and allowed to adhere before experimental treatment. Before steroid exposure, culture medium was replaced with medium containing charcoal-stripped serum. Cells were treated with vehicle, cortisol or 20*α*-dihydrocortisol at 500 nM for 20 min or 6 h as indicated. The primary-cell workflow is illustrated in Extended Data Fig. 8.

### RNA extraction, cDNA synthesis and qPCR

Cells in 12-well plates were lysed in 400 *µ*l TRI Reagent. After chloroform extraction, aqueous RNA was precipitated with isopropanol, washed in 75% ethanol, air-dried briefly and dissolved in RNase-free water. Samples were treated with RQ1 DNase-I and purified with the GeneJET RNA purification kit. RNA concentration was measured with Qubit. cDNA was synthesized with the qScript kit. SYBR Green qPCR was performed on a QuantStudio 6 Pro system. Expression was normalized to *β*-actin and calculated by the 2*^−^*^Δ^*^Cq^* method. Primer sequences for the *Nr4a3* loss-of-function experiments are listed in Supplementary Table 13.

### Protein extraction, immunoblotting and immunofluorescence

Cells were lysed on ice in 20 mM Tris-HCl, 145 mM NaCl, 5% glycerol, 1% Triton X-100 and protease inhibitor, pH 7.8. Lysates were cleared at 8,000*×g* for 10 min and protein concentration was measured with Bradford reagent. For immunoblotting, 50 *µ*g total protein was resolved on 12% SDS–PAGE and transferred to nitrocellulose with a Trans-Blot Turbo system. Membranes were blocked with 1% non-fat milk or 1% BSA in TBST, incubated with primary antibodies against NR4A3, pERK, ERK or *β*-actin overnight at 4 °C, followed by HRP-conjugated secondary antibodies. Chemiluminescence was recorded on an iBright CL750 and densitometry performed with iBright Analysis Software v5.4.0.

For immunofluorescence, cells were washed with PBS, fixed with 4% paraformaldehyde for 20 min, permeabilized with 0.1% Triton X-100 on ice for 20 min and blocked in 1% goat serum/1% BSA. CDX2, pERK or NR4A3 primary antibodies were applied at 1:50, followed by Alexa Fluor 488 secondary antibody at 1:100. Nuclei were counterstained with Hoechst. Images were acquired with a ZOE Fluorescent Cell Imager. Nuclear localization was quantified from colocalization of the protein signal with the nuclear counterstain. Representative pERK and NR4A3 fields are shown in Extended Data Figs. 8 and 9.

### *Nr4a3* siRNA knockdown

Primary epithelial cells at 40–50% confluence were transfected in 12-well plates with 40 pmol per well of mouse *Nr4a3* siRNA (ThermoFisher AM16708) or control siRNA (AM4611) using Lipofectamine 2000. Lipofectamine and siRNA were separately diluted in Opti-MEM, incubated for 5 min, combined and incubated for 20 min; 100 *µ*l complex was then added per well. After 48 h, cells received vehicle or 500 nM 20*α*-dihydrocortisol for 6 h before RNA and protein collection. Knockdown was evaluated by qPCR, immunoblotting and NR4A3 immunofluorescence.

### RNA sequencing and pathway analysis

RNA from mouse intestinal tissue/scrapings and primary epithelial cells was quality-assessed on a Fragment Analyzer. Individually barcoded poly(A)-selected RNA-seq libraries were prepared with the Kapa HyperPrep mRNA kit after rRNA depletion, using unique dual indexes. Libraries were quantified with Qubit and qPCR and sequenced on an Illumina NovaSeq 6000. Mouse intestinal libraries were sequenced as 150-nt paired-end reads; epithelial-cell libraries were sequenced as 100-nt single-end reads. Fastq files were generated with bcl2fastq v2.20 and quality assessed with FastQC. SeqKit summarized read statistics [23]; Trimmomatic removed adaptors and low-quality reads [24]; SortMeRNA removed residual rRNA [25]; and Salmon quantified transcript abundance [26]. Counts were filtered and normalized with edgeR, differential expression was analysed with limma, and KEGG enrichment was performed with clusterProfiler [27–29]. Multiplicity corrections are specified with the relevant analyses.

### Human study participants

Adults aged 45–75 years who self-identified as Black or White were recruited during screening or surveillance colonoscopy at Rush University Medical Center and the University of Illinois Chicago. Exclusion criteria included current malignancy or malignancy within the previous 5 years, history of colorectal cancer, alcoholism or illicit-drug use, gastrointestinal illness, significant medical conditions, therapeutic or vegetarian diets, anticoagulant use, medications interfering with digestion, antibiotics within 2 months, regular prebiotic/probiotic supplementation, pregnancy or nursing, and a menstrual period within the previous 6 months. The studies were approved by the institutional review boards of Rush University Medical Center (13102201), UIC/UIUC (2016-0495) and Purdue University (2022-1626). All participants provided written informed consent. Recruitment, exclusions and assay-specific sample sets are summarized in Extended Data Fig. 13 and Supplementary Tables 3–6.

Body weight was measured in duplicate to the nearest 0.1 kg on a calibrated Tanita BWB-800 scale and height was measured in duplicate to the nearest 0.1 cm with a stadiometer. BMI was calculated as kg m*^−^*^2^. Participants completed sociodemographic and tobacco-use questionnaires.

### Hair cortisol

Hair was collected from the crown vertex approximately 30 days after colonoscopy. The proximal 3 cm, representing approximately 3 months of growth, was used as an integrated measure of recent cortisol exposure [30–32]. Samples were weighed, washed twice with isopropanol, air-dried, ground, and extracted in methanol. Dried extracts were reconstituted in assay buffer and measured with the DetectX Cortisol enzyme immunoassay (Arbor Assays, K003-H1W/H5W) at the Center for Neuroendocrine Studies. Participants reporting corticosteroid-product use and values deemed biologically implausible (*≤* 0 or *≥* 750 pg mg*^−^*^1^) or more than three standard deviations from the mean were excluded according to the prespecified quality-control rules [33].

### Human faecal metagenomic sequencing and profiling

Stool was collected approximately 30 days after colonoscopy. DNA was extracted with the QIAGEN DNeasy PowerSoil kit and sent on dry ice to Monash University. Shotgun libraries were sequenced as paired-end 150-bp reads. Reads were quality-trimmed and depleted of human hg38 sequence as described previously [34]; remaining read pairs were interleaved with seqtk mergepe v1.5. Taxonomic and functional profiles were generated with MetaPhlAn 4 and HUMAnN v3.9 using mpa_vJun23_CHOCOPhlAnSGB_202403 [35, 36]. Species-level MetaPhlAn outputs were merged across samples for diversity and abundance analyses.

### Human faecal *desC* qPCR and steroid analysis

Faecal *desC* was quantified with SYBR Green qPCR on a QuantStudio 6 Pro. *desC*-specific primers were analysed alongside universal 16S rRNA gene primers (Supplementary Table 15), and relative abundance was calculated by 2*^−^*^Δ^*^Cq^*.

For steroid measurements, stool collected approximately four weeks after colonoscopy was stored at *−*80 *^◦^*C, lyophilized, homogenized by ultrasonication and extracted with methanol for 24 h at *−*20 *^◦^*C. Supernatants were collected after centrifugation, dried under nitrogen and stored at *−*80 *^◦^*C. Residues were dissolved in 20% methanol, spiked with internal standard and analysed for 20*α*-dihydrocortisol on the Agilent 6495A LC–MS/MS platform used for mouse samples.

### Geospatial exposures

Residential addresses were geocoded and linked to census-tract/community-area measures. Homicide rates per 100,000 residents and percentage poverty were obtained from the Chicago Health Atlas for 2000–2018 and averaged across the available years [37]. Self-identified race is used here to index social and environmental context and is not treated as a biological category.

## Statistical analysis

Microbiome analyses were performed in R v4.4.3. Species richness was calculated with vegan v2.6-10 and compared between groups by two-sided Wilcoxon rank-sum test. Bray–Curtis dissimilarity was calculated from species-level relative abundances and tested by PERMANOVA with 999 permutations; principal-coordinate analysis was used for visualization. *C. scindens* relative abundance was compared by Wilcoxon rank-sum test.

Normality of faecal *desC*, hair-cortisol and mouse metabolite measurements was assessed with the Shapiro–Wilk test. Nonparametric two-group comparisons used Wilcoxon rank-sum tests; families of metabolite comparisons were adjusted with the Benjamini–Hochberg procedure. Gene and protein measurements were compared by two-sided *t*-tests with the multiplicity correction specified in the corresponding analysis. Pathway-enrichment *P* values were adjusted for false discovery rate. For human faecal 20*α*-dihydrocortisol, in which many values were below the detection limit, detection frequency was first compared by Fisher’s exact test and concentrations were then compared by Wilcoxon rank-sum test only among samples with detectable metabolite. Exact sample sizes, definitions of centre/error bars and corrections are given in the figure legends or Supplementary Tables.

